# Macrophages tune responses to pathogen dynamics through TLR4 stimulation memory and by licensing susceptibility to IL-10

**DOI:** 10.1101/2024.03.28.587272

**Authors:** Hannes Bongartz, Christopher T. Boughter, Bernadette Marrero, Thorsten Prüstel, Clinton Bradfield, Julia Gross, Rachel A. Gottschalk, Aleksandra Nita-Lazar, Iain D. C. Fraser, Martin Meier-Schellersheim

## Abstract

Macrophages must scale inflammation to the trajectories of infections, escalating as pathogen loads rise, resolving as they fall. The cytokine IL-10 and its chromatin-level effector BCL-3 are critically important anti-inflammatory agents, yet how they avoid muting newly recruited cells before those cells read their own pathogen input has been unclear. Applying systematically varied consecutive TLR4 (Kdo2-Lipid A) stimuli to bone marrow-derived macrophages, we identify two coupled features. First, macrophages retain a quantitative memory of prior stimulation: only when a secondary TLR stimulus matches or exceeds a prior one, IκBα degradation, NF-κB/MAPK activation, and cytokine output increase. Second, IL-10 susceptibility is itself gated by TLR4 history: even a 100-fold IL-10 excess fails to suppress TNF-α in weakly primed cells. Both trace to history- and IL-10-dependent BCL-3 recruitment with p65 displacement at the Tnf κB site. Modeling shows how this licensing logic enables trajectory-aware responses that clear pathogens while limiting tissue damage.

## Introduction

Macrophages, together with other innate immune cells, form the primary defense against many pathogens. They sense pathogen-associated molecular patterns through pattern-recognition receptors such as Toll-like receptor 4 (TLR4), which mediates the response to lipopolysaccharide of gram-negative bacteria and activates the MyD88-dependent mitogen-activated protein kinase (MAPK) and nuclear factor-κB (NF-κB) cascades as well as the TRIF/IRF interferon arm, culminating in the production of pro- and anti-inflammatory cytokines (*1–5*). Pathogen loads, however, are not static. They rise during the establishment of an infection, plateau, and fall as the pathogen is contained or cleared, and the response that is appropriate at a given moment depends on this trajectory: a cell facing a given current input faces a very different task if the load is still increasing than if it is already being controlled. Sustained inflammation may unnecessarily induce tissue damage when deployed against an already-resolving threat, while premature de-escalation against a still rising load risks failure of pathogen control. Innate immune cells must therefore not merely detect pathogens but also assess whether their exposure is rising or declining.

That macrophages do not respond identically to every TLR4 stimulus has been documented for decades. Low-dose pre-exposure can sensitize cells to subsequent challenges (*6–10*), while strong or prolonged stimulation produces a hypo-responsive, tolerant state (*11–15*). A recent systematic study profiled single-cell NF-κB activation dynamics across numerous pairwise combinations of inflammatory ligands in primary macrophages and showed that the dose and duration of prior stimulation are encoded in subsequent NF-κB signaling, with both tolerance and priming arising in ligand- and dose-dependent ways (*10*). That work, however, varied only the strength of the primary stimulus against a single secondary dose, and focused primarily on combinations of different TLR ligands. Whether the amplitude of a secondary response within the same TLR pathway is set by a quantitative comparison between secondary and primary stimulus strengths has not been resolved. Further, it has remained unexplored whether such a comparison extends from NF-κB dynamics to MAP-kinase activation, IκBα degradation, and downstream cytokine and chemokine output.

At the molecular level, the components of the cellular signaling processes that dampen macrophage activation are well characterized. The anti-inflammatory cytokine IL-10, signaling through STAT3, is a central node (*15–29*), and the IL-10-inducible atypical IκB family member BCL-3 is an established chromatin-level mediator of cytokine gene suppression (*30, 31*). What has remained unclear is how anti-inflammatory regulation could preserve the system’s ability to follow the trajectory of an infection. If IL-10 acted on every cell simply as a function of its concentration, then once the early responders in an infected tissue had begun secreting IL-10, every macrophage newly recruited to the site would be immediately conditioned by the ambient IL-10 of its neighbors, muted before it could read its own pathogen input. The same problem arises from IL-10 contributed by other sources: regulatory T cells, B cells, neighboring leukocytes and, at mucosal sites, epithelial- and microbiota-driven signals supply ambient IL-10 at steady state (*32, 33*). Moreover, several intracellular bacteria and protozoa induce host IL-10 (*34*) while certain viruses encode IL-10 mimics that engage the host receptor as an immune-evasion strategy (*35, 36*). Under a concentration-only model, the collective effect of such inputs would be to collapse the trend information that distinguishes a rising load from a falling one into a uniform suppressed state, as opposed to permitting a graded, trajectory-aware response.

Two coupled questions follow that previous work has not fully addressed. First, do macrophages register the strength of prior TLR4 stimulation quantitatively, in a form that lets each cell compare a current input to its own history rather than respond only to its absolute current level? Second, is a cell’s susceptibility to IL-10 itself a fixed property or is it conditioned by its TLR4 history, such that cells which have not yet committed to a TLR4 response remain unaffected by ambient IL-10? Whereas many studies have elucidated the anti-inflammatory action of IL-10 and of its influence on the chromatin level functions of BCL-3, they have largely used uniform strong TLR4 stimulation (*16, 23, 26, 31, 37, 38*) and have not performed the dose-resolved comparisons required to answer these questions.

In this study, we systematically varied consecutive stimulations of BMDM with the TLR4 ligand Kdo2-Lipid A (KLA), spanning a wide range of doses and dose combinations in primary and secondary stimuli, and titrated exogenous IL-10 against both weak and strong primary stimulation. We find that BMDM register the strength of prior TLR4 input quantitatively: secondary stimuli amplify activation of MAPK, NF-κB and IκBα degradation and increase pro-inflammatory cytokine and chemokine output only when their strengths match or exceeds the primary stimuli, encoding the direction of a temporal trend rather than its instantaneous level. We further find that the susceptibility of BMDM to IL-10 is itself gated by TLR4 history, with even an approximately 100-fold excess of exogenous IL-10 failing to suppress TNF-α production in cells that have not received strong prior stimulation. We trace both features to a chromatin-level mechanism at the Tnf κB site, where TLR4 history and IL-10 jointly govern BCL-3 recruitment with reciprocal displacement of p65, in a manner controlled by the chromatin-binding competence of p50/BCL-3 rather than its abundance or nuclear localization. The result is a coincidence-detector logic that, rather than using them as indiscriminate suppressors, lets the anti-inflammatory components IL-10 and BCL-3 function as part of a regulator reading the cells stimulation histories.

Intuitively, such a cell-intrinsic logic may allow a macrophage population to stay aligned with the trajectory of an infection while indiscriminate suppression of innate function by ambient IL-10 may lead to a failure of pathogen control. To test this intuition quantitatively and explore whether history-gated IL-10 susceptibility may act as a discriminatory suppressor at the population level, we built a simple computational model of macrophages responding to an evolving infection and compared three regulatory logics: (i) history-gated licensing combined with quantitative memory, (ii) suppression set only by the ambient IL-10 concentration, and (iii) no IL-10 regulation of macrophage responses. Robust modeling results indicate that the licensing logic allows the macrophage population to control the pathogen while limiting inflammation-driven tissue damage.

## Results

### TLR4-ligand induced signaling responses show sensitization, adaptation and quantitative memory of primary stimulation

To examine how the TLR4 pathway balances pro- and anti-inflammatory output when stimulation strength changes over time, we exposed BMDM to consecutive KLA stimuli applied as systematically varied concentration sequences (Fig. 1A). Because surface TLR4 levels shape the balance between MyD88- and TRIF-dependent signaling (*14*), we first asked how surface TLR4 changes with KLA dose, extending earlier data limited to single LPS doses or to ≤90 min duration (*12, 39*). A 4 h primary challenge with 0, 1, 10 or 100 nM KLA reduced surface TLR4 in a dose-dependent manner, leaving ∼10% of receptors after 100 nM (Fig. 1B).

**Fig. 1:**
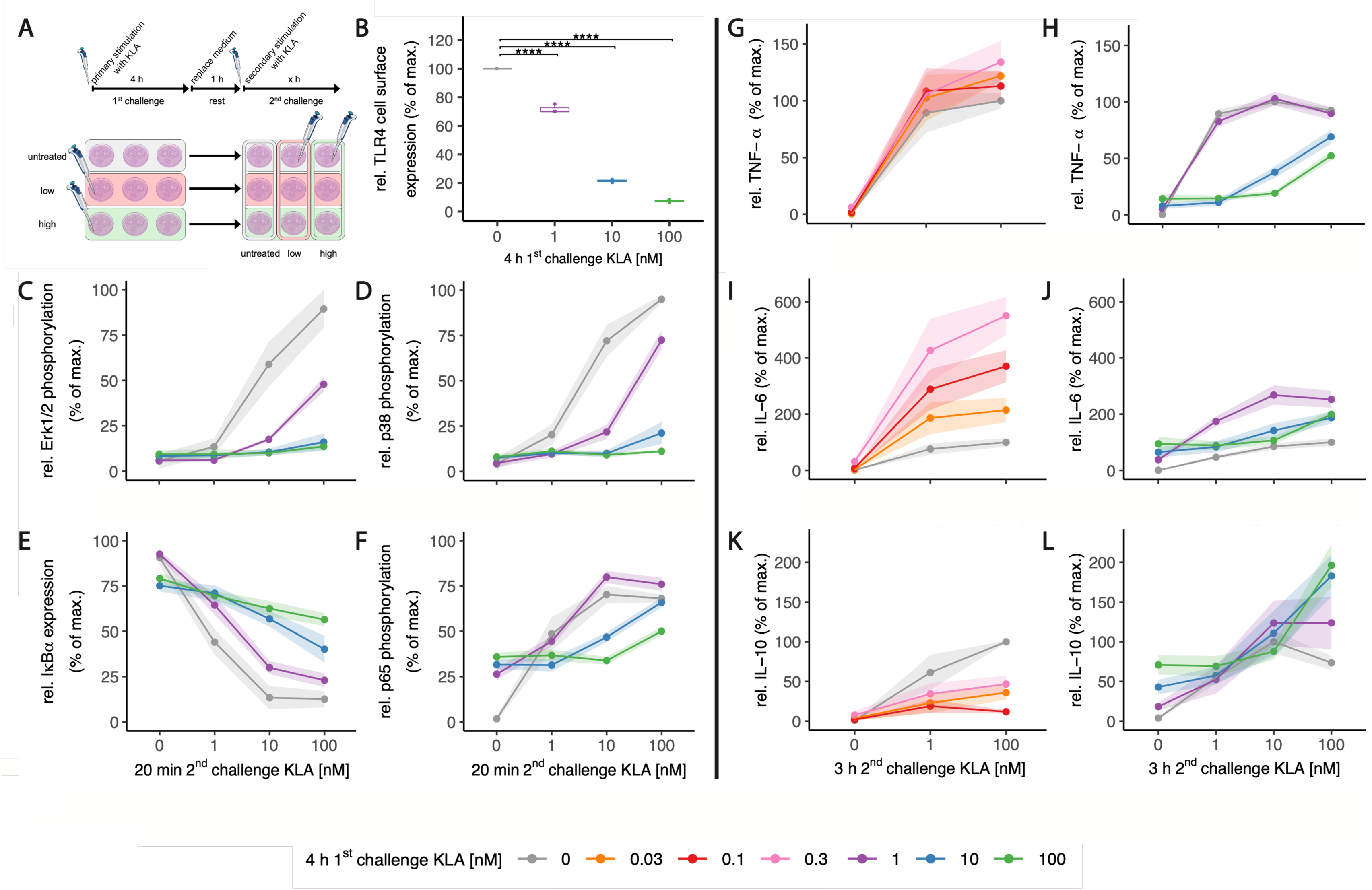
**(A) Treatment regimen of 1^st^ challenge and 2^nd^ challenge with KLA of primary murine macrophages.** Bone marrow derived macrophages (BMDMs) were differentiated from isolated bone marrow cells of wildtype C57BL/6J mice by M-CSF. Prior to experiment, BMDMs were incubated with 5% FBS DMEM without M-CSF. BMDMs were treated with M varying doses of KLA (1^st^ challenge) for 4 hrs. Then, after washing and replacing the medium, cells were rested for 1 hr and were restimulated (2^nd^ challenge) with N varying doses of KLA resembling a two-stage dosing design yielding an NxM matrix of dose combinations. Depending on the experimental read-out, 2^nd^ challenge times differ. **(B) TLR4 internalization becomes stronger as primary KLA concentrations increase.** During 1^st^ challenge, cells were stimulated with 0, 1, 10 or 100 nM KLA for 4 hrs. Cells were washed, and medium was replaced with 5% FBS DMEM and were incubated for 1 hr. Subsequently, cells were stained for TLR4 surface expression. Surface expression was determined by using a LSRII flow cytometer (BDBiosciences). Raw data was processed with FlowJo™. Median fluorescence intensities were corrected to the median fluorescence intensity of the TLR4 detecting antibody from a TLR4-KO cell line (unspecific binding background), normalized to untreated BMDMs, and represented in percent of untreated control sample (grey, set as 100%). Data include 3 replicates (n=3). Each replicate was performed with BMDMs from different mice. P-values: Kruskal–Wallis test with post-hoc Dunn-Bonferroni comparisons of stimulated samples with untreated sample (planned comparisons). **(C-F) Dose dependent activation and adaptation of MAP kinases, IkBa and p65.** During 1^st^ challenge, cells were stimulated with 0, 1, 10 or 100 nM KLA. Restimulation was performed using 0, 1, 10 or 100 nM KLA. After 20 mins, cells were intracellularly stained for (C) phosphorylated Erk1/2, (D) phosphorylated p38, (E) IkBa, and (F) phosphorylated p65. Median fluorescence intensity of samples was determined by using a LSRII flow cytometer (BDBiosciences). Raw data was processed with FlowJo™. Data is given in % of maximal median fluorescence intensity within each replicate (set as 100%) and was normalized by subtracting median fluorescence intensity of the sample with the detected minimal median fluorescence intensity (set as 0%). Each replicate was performed with BMDMs from different mice. Data include at least 4 replicates (n=4) and are shown as mean ± standard error of mean (SEM). **(G-L) KLA concentration during 1^st^ challenge dictates responsiveness of cytokine release in response to restimulation.** During primary and secondary stimulation, BMDMs were incubated with 5% FBS DMEM. During 1^st^ challenge, cells were stimulated with 0, 0.03, 0.1,0.3 nM KLA (G,I,K) or with 0, 1, 10 or 100 nM KLA (H,J,L). Restimulation was performed using 0, 1, 10 or 100 nM KLA. After 3 hrs, supernatants were collected, processed with the LegendPlex™ Multiplex Assay Kits and cytokine levels of (G,H) TNF-a, (I,J) IL-6, and (K,L) IL-10 were determined. Raw data was processed with LegendPlex™ Desktop software. Each replicate was performed with BMDMs from different mice. Data include 6 replicates (n=6). Data was normalized to the mean maximal cytokine secretion induced by restimulation in naïve (0 nM KLA primary challenge) cells (set as 100%). Data are shown as mean ± SEM.

To test whether macrophages with a significantly depleted surface TLR4 pool could still discriminate stimulus strength, we washed and rested the cells for 1 h after the 4 h primary challenge (“1st challenge”) and restimulated them for 20 min (“2nd challenge”), the time at which MAPK phosphorylation peaks for intermediate-to-strong signals (*8*). Increasing primary concentrations progressively weakened secondary MAPK responses and, notably, Erk1/2 and p38 were re-activated only by secondary stimuli exceeding the primary dose: for example, after 1 nM priming, both kinases responded only when the second stimulus surpassed 1 nM (Fig. 1C,D; target genes in Fig. S1A–C).

IκBα degradation and p65 phosphorylation were more readily induced by restimulation, responding when the secondary stimulus matched or exceeded the first and remaining robust even after strong priming (Fig. 1E,F), consistent with our earlier work (*8*) and indicating that NF-κB signaling stays responsive despite the loss of most surface TLR4.

We next asked whether this behavior propagated to downstream cytokine output. Low-dose priming (≤0.3 nM KLA) sensitized TNF-α and IL-6 responses to secondary challenge, increasingly so with higher primary dose (Fig. 1G,I), however past priming concentrations of 1 nM, this trend changed and secondary responses became progressively reduced (Fig. 1H,J). This was not a full shut-down: after 10 or 100 nM priming, TNF-α and IL-6 were still produced, but only in response to secondary stimuli reaching or exceeding the primary dose – mirroring p65 phosphorylation (Fig. 1F) and indicating that the cells respond selectively to stimuli that surpass their stimulation history. CXCL-1 behaved like TNF-α (Fig. S3C), while IFN-β was an exception, losing responsiveness sharply after priming with ≥1 nM KLA (Fig. S2D, S3D). In contrast to the pro-inflammatory mediators, IL-10 secretion depended non-monotonically on the stimulation sequence, with a minimum of restimulation-induced secretion at low-intermediate priming doses (0.03 −0.3 nM KLA) (Fig. 1K,L; Fig. S2E).

These findings suggested that IL-10-mediated regulation of TNF-α and IL-6 may itself depend on the history of TLR4 stimulation.

### Downstream responses show strongly varying impact of TRL4 stimulation history on IL-10 mediated anti-inflammatory effects

To identify which responses were directly controlled by IL-10, we applied a non-signaling IL-10-receptor-blocking antibody, or an isotype control, throughout primary and secondary stimulation. Proximal signaling was largely IL-10- independent: p38 phosphorylation showed no dependence on IL-10 under our conditions, in contrast to some earlier reports (*24*), and p65 phosphorylation was only slightly increased by IL-10R blockade after intermediate (1 nM) priming (Fig. 2A,B). Cytokine output, by contrast, was strongly affected. The negative regulation of IL-6 seen after intermediate and strong priming was IL-10-dependent across all restimulation doses (Fig. 2C). In contrast, TNF-α had shown a much stronger suppression following strong priming (10 or 100 nM) than for priming with up to 1 nM KLA (Fig. 1H) and this suppression was abolished by IL-10R blockade (Fig. 2D). The effect was again selective: CCL-4 and CCL-5 were largely IL-10-independent, whereas CXCL-10 and IL-12p70 increased upon IL-10R blockade, but only after priming with ≥0.3 nM KLA (Fig. S2, S3). IL-10 itself rose when its receptor was blocked, consistent with autocrine feedback control of its expression (*40, 41*).

**Fig. 2:**
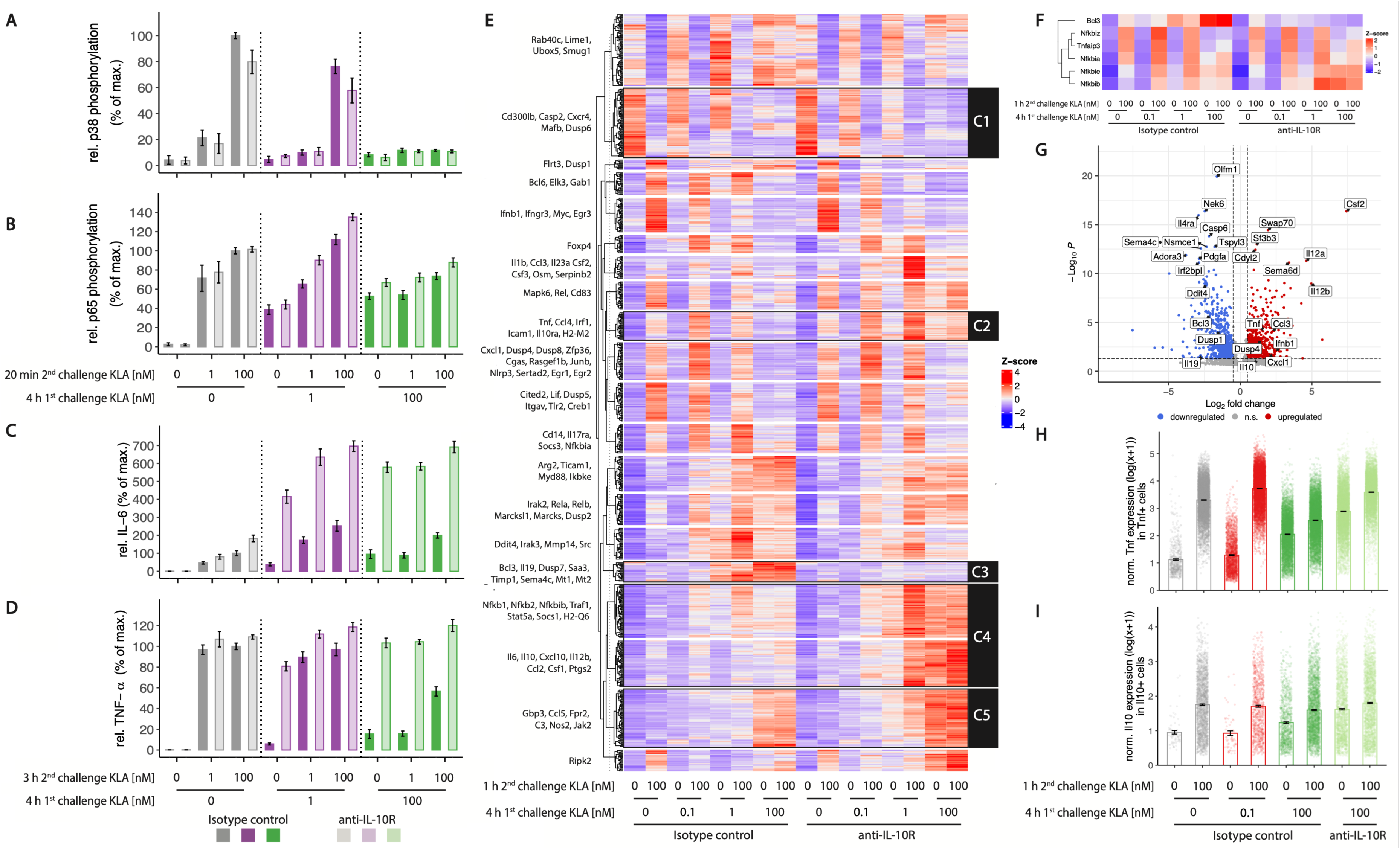
**(A-D) Blocking IL-10 signaling does not substantially affect upstream TLR4-ligand-induced MAPK and NFkB signaling but downstream cytokine secretion.** During primary and secondary stimulation, BMDMs were incubated with 5% FBS DMEM containing either IL-10R blocking antibody (weakly saturated bars) or its isotype control (saturated bars). During 1^st^ challenge, cells were stimulated with 0, 1, or 100 nM KLA. Restimulation was performed using 0, 1, or 100 nM KLA. (A,B) After 20 mins restimulation, cells were intracellularly stained for (A) phosphorylated p38, and (B) phosphorylated p65. Median fluorescence intensity of samples was determined by using a LSRII flow cytometer (BDBiosciences). Raw data was processed with FlowJo™. Each replicate was performed with BMDMs from different mice. Data include 4 replicates (n=4), and was normalized to the maximal median fluorescence intensity induced by restimulation in naïve (0 nM KLA primary challenge) cells (set as 100%). (C,D) After 3 hrs restimulation, supernatants were collected, processed with the LegendPlex™ Multiplex Assay Kits and cytokine levels of (C) IL-6, and (D) TNF-α, were determined. Raw data was processed with LegendPlex™ Desktop software. Each replicate was performed with BMDMs from different mice. Data include 6 replicates (n=6, see Fig. S3,S4), and was normalized to the mean maximal cytokine secretion induced by restimulation in naïve (0 nM KLA primary challenge) cells (set as 100%). Data are given as mean ± SEM. **(E-G) IL-10 mediates hypo-responsiveness of pro-inflammatory target genes at transcriptional level.** During primary and secondary stimulation, BMDMs were incubated with 5% FBS DMEM containing either IL-10R blocking antibody or its isotype control. During 1^st^ challenge, cells were stimulated with 0, 0.1, 1 or 100 nM KLA. Restimulation was performed using 0 or 100 nM KLA. After 1 hr, total RNA was isolated. (E) RNA from each sample was sequenced and analyzed. Expressed RNA significantly changed by TLR4 activation was clustered according to row-wise Z-scores which were derived from DESeq2-normalized counts, and representative genes for each cluster as well as their differential sensitivity towards primary stimulation and IL-10 were annotated. (F) Gene-expression of several NFkB pathway feedback inhibitors. (G) Differentially expressed gene analysis comparing effect of IL-10 receptor block (without vs. with anti-IL-10R) in 100 nM KLA primary and 100 nM KLA secondary stimulated cells identified IL-10 dependent genes. **(H,I) Gene expression levels in single cells together with size of gene expressing subpopulation determine the TLR4 activation history dependence of pro- and anti-inflammatory response.** During primary and secondary challenge, BMDMs were incubated with 5% FBS DMEM containing either IL-10R blocking antibody or its isotype control. During primary stimulation, cells were treated with 0, 0.1, or 100 nM KLA. Restimulation was performed using 0 or 100 nM KLA. After 1 hr, cells were subjected to single cell RNA sequencing. (H) Normalized Tnf expression in Tnf positive cells. (I) Normalized Il10 expression in Il10 positive cells. Data shown from 2 independent replicates.

These findings suggested that there are complex patterns of gene expression control related to the strength of TLR4 signaling and to IL-10 exposure. To better characterize history-dependent responses and look for coherent clusters of regulated genes at a transcriptome-wide level, we performed RNA-seq on BMDM primed with 0, 0.1, 1 or 100 nM KLA and restimulated with 0 or 100 nM, with or without IL-10R blockade.

A 60 min challenge of naïve cells significantly altered 776 genes (Fold of control ≥ 2; Fig. 2E), which were grouped into multiple clusters, among which five were particularly distinguished by their dependence on KLA dose and on IL 10 (Table 1; full cluster gene lists in Table S1).

**Table 1.** Gene-expression clusters defined by TLR4 dose and IL-10 dependence.

| Cluster | Dependence on KLA dose | Influence of IL-10 | Representative genes |
| --- | --- | --- | --- |
| C1 | Highest in resting cells, decreases with increasing KLA | Mostly IL-10 independent | Cd300lb, Casp2, Cxcr4, Mafb, Dusp6 |
| C2 | Induced by KLA; adapts to high primary dose | Adaptation requires IL-10 | Tnf, Ccl4, Irf1, Icam1, Il10ra, H2-M2 |
| C3 | Induced by KLA; strongest after high primary dose | Induction requires IL-10 | Bcl3, Il19, Dusp7, Sema4c, Timp1, Mt1, Mt2 |
| C4 | Induced by strong KLA | Strongly suppressed by IL-10 | Nfkb1, Nfkb2, Nfkbib, Il6, Il12b, Il27, Cxcl10, Socs, IL10, IL12b, Ccl2 |
| C5 | Increases with KLA dose | Less strongly suppressed by IL-10 | Gbp3, Nos2, Ccl5, Jak2, Fpr2, C3 |

The clusters most relevant to adaptation were C2 and C3. C2 comprised pro-inflammatory genes (e.g. Tnf, Ccl4) that show high expression in response to primary stimulation across the dose range used, whereas they exhibit strong negative regulation after high primary KLA stimulation. The latter adaptation is lost upon IL-10R blockade. Notably, the IL-10 receptor itself fell into C2, indicating that strong pro-inflammatory signaling also raises the cells’ capacity to sense IL-10. C3 comprised IL-10-dependent anti-inflammatory genes (e.g. Bcl3, Il19, Dusp7) most strongly induced after 100 nM priming, many associated with the M2 phenotype (*42–45*).

Some other established negative regulators of TLR4 signaling, such as several members of the NFκB family exhibited patterns of TLR4 stimulation dependencies (Fig. 2F) that were different from the behavior of BCL-3.

Cluster C1, highly expressed in resting cells and progressively lost with increasing KLA, independently of IL-10 (Fig. 2E), includes genes of a tissue-resident identity program (e.g. Mafb, Cxcr4) dismantled during inflammatory activation, while clusters C4 and C5 contained strongly KLA-driven, M1-associated genes differing chiefly in their degree of IL-10 sensitivity. The mRNA profiles of most cytokines and chemokines recapitulated their secretion (compare Fig. S4 with Fig. 1, S2, S3), indicating that the observed regulation is largely transcriptional.

**Fig. 3:**
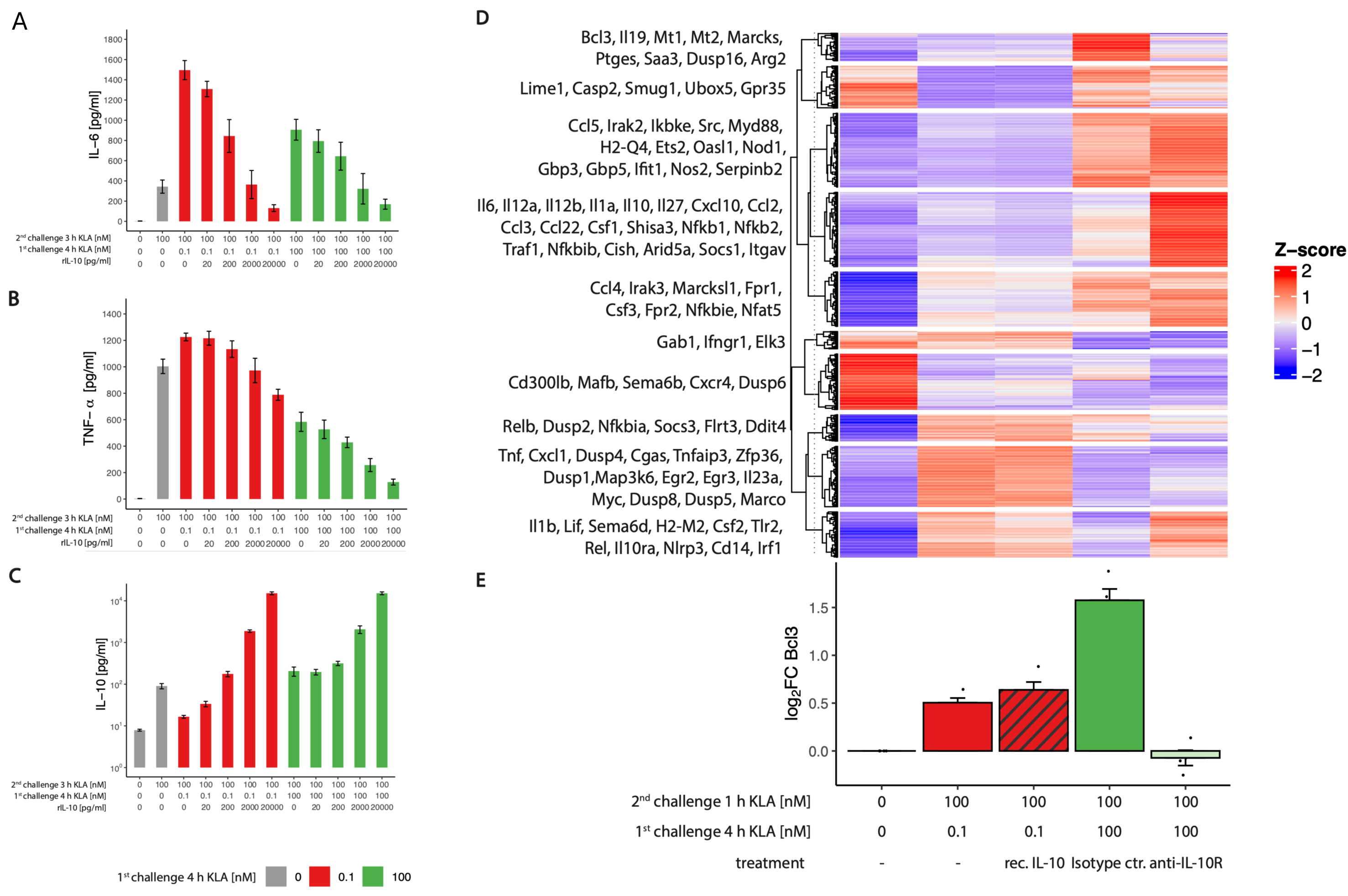
Broad induction of hypo-responsiveness requires prior strong TLR4 simulation, not just IL-10. (A-C) During primary stimulation, cells were treated with 0, 0.1, or 100 nM KLA. Restimulation was performed using 0 or 100 nM KLA. 20 min after each KLA stimulation, recombinant murine IL-10 was added in depicted concentrations. After 3 hrs restimulation, supernatants were collected, processed with the LegendPlex™ Multiplex Assay Kits and cytokine levels of (A) IL-6, (B) TNF-α, and (C) IL-10 were determined. Raw data was processed with LegendPlex™ Desktop software. Each replicate was performed with BMDMs from different mice. Data include 9 replicates (n=9) and are given as mean ± SEM. (D,E) During primary and secondary challenge, BMDMs were incubated with 5% FBS DMEM containing either IL-10R blocking antibody or its isotype control. During primary stimulation, cells were stimulated with 0, 0.1, or 100 nM KLA. Restimulation was performed using 0 or 100 nM KLA. 20 min after each KLA stimulation, 200 pg/ml of recombinant murine IL-10 was added. After 1 hr restimulation, total RNA was isolated, sequenced and analyzed. (D) Expressed RNA significantly changed by TLR4 activation was clustered according to row-wise Z-scores which were derived from DESeq2-normalized counts. (E) Log₂ fold changes for Bcl3 mRNA expression were calculated from DESeq2-normalized counts as mean expression ratios versus control (pseudocount = 1). Data are given in mean ± SEM of n=4 replicates.

Prior work had shown that population responses of IL-10 were regulated by the fraction of IL-10 producers whereas single-cell TNF-a production may vary depending on the TLR4 stimulus the cells received (*9, 46, 47*). We therefore analyzed how TLR4 stimulation history shaped single cell expression levels of TNF-α and IL-10. The 0.1/100 protocol generated more high-Tnf cells than 100/100 and closely resembled 100/100 with the IL-10 receptor blocked (Fig. 2H). The history dependence of TNF-α output is therefore encoded in per-cell expression intensity rather than in the size of the producing population, resolving the apparent discrepancy with the secretion data (Fig. 1G,H). Il10 showed the converse: per-cell expression was largely invariant across protocols (Fig. 2I, Fig. S7), so that total IL-10 output was set by the fraction of producing cells (*9, 46, 47*).

### Susceptibility to IL-10 requires strong prior TLR4 stimulation, not merely IL-10 itself

In contrast to the behavior of IL-6 (Fig. 2C), the IL-10-dependent hypo-responsiveness of TNF-α occurred only in macrophages pre-exposed to the highest KLA concentrations (Figs. 2D, S2, S3). One potential explanation was that suppressing TNF-α requires more IL-10 than suppressing IL-6 – which is directly controlled by IL-10/STAT3 – so that only strong TLR4 stimulation generates sufficient IL-10 (*46*). To test this, we primed BMDM for four hours with a low (0.1 nM) or high (100 nM) KLA dose, restimulated for three hours with 100 nM KLA, and titrated recombinant IL-10 from 20 to 20,000 pg/ml, supplied during both, primary and secondary stimulation. Importantly, 200 pg/ml corresponded to the maximum IL-10 produced by the cells themselves after 100 nM KLA (Fig. S3 E) and was sufficient to suppress pro-inflammatory output after strong priming with 10 or 100 nM KLA (Fig. S3 A, B, C).

Consistent with an unconditional susceptibility towards suppression by high concentrations of IL-10, IL-6 responses were suppressed below the level seen after strong TLR4 priming (Fig. 3A) by IL-10 concentrations above 200 pg/ml.

Confirming that the role of IL-10 for the regulation of TNF-α responses is quite different from its role for IL-6, even the highest concentrations of exogenous IL-10 failed to suppress TNF-α production to the level reached after strong priming (compare rightmost red column to leftmost green column in Fig. 3B). This suggests that the divergent behavior of IL-6 and TNF-α does not simply reflect different sensitivities to IL-10.

The responses of other cytokines and chemokines towards combinations of weak or strong TLR4 stimulations and exogenous IL-10 were highly heterogenous (Fig. S5). Interestingly, high concentrations of recombinant IL-10 could strongly suppress IFN-β and CXCL-1, however not beyond the suppression seen for strong TLR4 stimulation (Fig. S5 A, B).

In contrast, CCL-2 and CXCL-10 were only weakly secreted after low-dose pre-stimulation and showed only limited susceptibility to exogenous IL-10, while strongly pre-stimulated cells secreted higher levels of these chemokines and became responsive to IL-10-mediated suppression (Fig. S5C,F).

Macrophages pre-stimulated with 0.1 nM KLA and subsequently treated with 200 pg/ml exogenous IL-10 exhibited reduced IL-10-induced STAT3 phosphorylation compared with naïve cells treated with the same concentration of IL-10 (Fig. S5H). This suggests that susceptibility of macrophages towards the anti-inflammatory IL-10 may be determined by their TLR4 activation history in a target-gene specific manner.

Transcriptome-wide, the difference between the behavior of Il6 and that of other genes was striking. Reminiscent of what we had observed for TNF-α and IL-6 production (Fig. 3A, B), 200 pg/ml recombinant IL-10, although sufficient to suppress IL-6, altered only a small subset of genes in low-dose-primed macrophages (Fig. 3D), with the Il6-containing cluster the only one whose behavior approached that of strong priming. Direct suppression of IL-6 thus appears to be the exception rather than a model for the general action of IL-10 as anti-inflammatory agent. Consistent with this, the Bcl3-containing cluster showed almost no response to exogenous IL-10 when cells had received only weak priming, whereas dual strong stimulation strongly upregulated Bcl3 in an IL-10-dependent manner (Fig. 3E). Although very high IL-10 concentrations were found to induce BCL-3 previously (*31*), the highest IL-10 levels produced in our assays, while sufficient to suppress IL-6, did not induce Bcl3 without prior strong TLR4 stimulation. Post-transcriptional control may also contribute: Zfp36, encoding the TNF-mRNA-destabilizing protein TTP, depended strongly on TLR4 history but only weakly on IL-10 (Fig. 2E, 3D), in line with prior reports (*40, 48*).

### TLR4 history and IL-10 condition the chromatin association of BCL-3 independently of its nuclear abundance

Because BCL-3 contributes to transcriptional repression by competing with p65 for p50-mediated κB-site binding (*30, 37, 38, 49*), we first asked whether the nuclear abundance of BCL-3, p50 and p65 tracked target-gene regulation. Restimulation-induced nuclear accumulation of p50 and p65 increased with stronger secondary KLA but ceased to increase after 1 and 100 nM priming (Fig. 4A,B; Fig. S6B,C,E,F), consistent with upstream IκBα degradation and p65 phosphorylation (Fig. 1E,F). BCL-3 was predominantly nuclear (Fig. S6G), but its nuclear accumulation decreased with higher primary doses (Fig. 4C). IL-10R blockade only slightly reduced the nuclear levels of all three factors (Fig. S6B–D). Neither nuclear accumulation nor mRNA levels of BCL-3, p50 or p65 therefore predicted target-gene suppression, arguing against a simple mass-action competition and indicating that nuclear NF-κB translocation cannot be used to quantify the strength of upstream receptor stimulation.

**Fig. 4:**
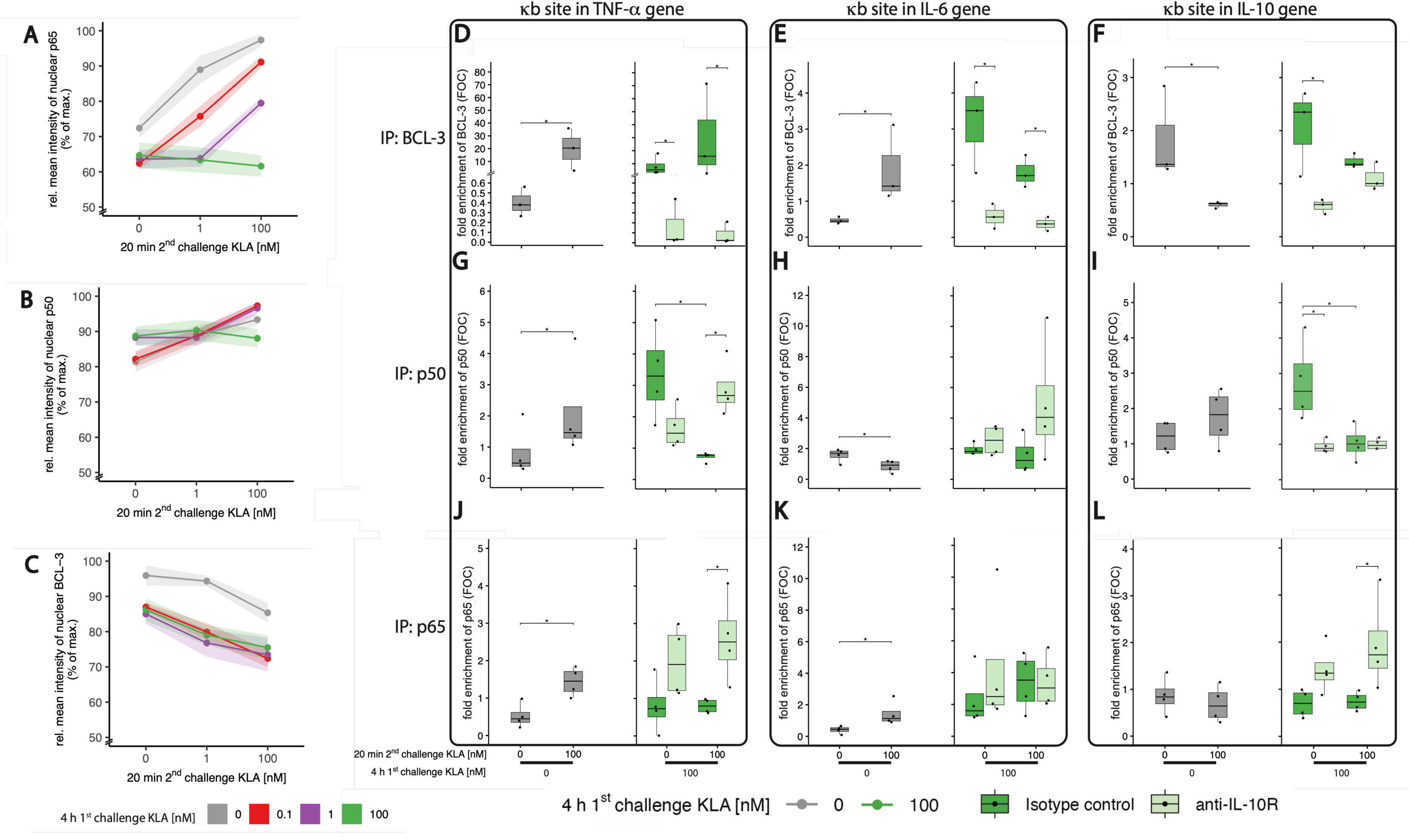
TLR4 primary and secondary stimulation as well as Il-10 signaling affect nuclear localization and binding of BCL-3, p65 and p50 at kB sites of target genes. (A-C) During primary and secondary stimulation, BMDMs were incubated with 5% FBS DMEM containing either IL-10R blocking antibody or its isotype control. During 1^st^ challenge, cells were stimulated with 0, 0.1, 1 or 100 nM KLA. Restimulation was performed using 0, 1 or 100 nM KLA. After 20 mins, cells were intracellularly stained for (A) p65, (B) p50, and (C) BCL-3 expression and localization was determined by using a CellInsight CX7 Pro HCS Platform (Thermo Fisher Scientific) equipped with an 40x objective lens. Raw data was processed with CellProfiler™. Data is given in % of maximal median fluorescence intensity within each replicate (set as 100%). Each replicate was performed with BMDMs from different mice. Data include at least 6 replicates (n=6) and are given as mean ± standard error of mean (SEM). (D-L) During primary and secondary challenge, BMDMs were incubated with 5% FBS DMEM containing either IL-10R blocking antibody or its isotype control. During 1^st^ challenge, cells were stimulated with 0 or 100 nM KLA. Restimulation was performed using 0 or 100 nM KLA. After 20 mins, cells were fixed to cross-link chromatin and protein interactions and chromatin was sheared. Chromatin immuno-precipitation (ChIP) of either, cross-linked (D-F) BCL-3, (G-I) p50 or (J-L) p65 – chromatin complexes was conducted with BCL-3, p50 or p65 targeting antibodies and, in parallel, with corresponding Isotype control antibodies. DNA from purified Protein – chromatin complexes was quantified with qPCR using specific primers for kB sites in TNF-a (D,G,J), IL-6 (E,H,K) and IL-10 (F,I,L) genes. The amount of chromatin DNA in anti-BCL-3, anti-p50 and anti-p65 antibody immunoprecipitated samples was normalized to DNA amounts in their respective Isotype-control antibody immuno-precipitated samples (fold enrichment). Each replicate was performed with BMDMs from different mice and data include at least 3 replicates (n=3). P-values: Wilcoxon test for KLA stimulation in naïve BMDMs (grey) and Kruskal–Wallis test with post-hoc Dunn- Bonferroni comparisons for 100 nM KLA pre-exposed BMDMs (± anti-IL-10R, green and light-green).

We therefore examined factor occupancy directly by chromatin immunoprecipitation followed by qPCR at known κB sites in the TNF-α, IL-6 and IL-10 genes. In naïve cells, a 20 min 100 nM challenge enriched BCL-3 at the TNF-α and IL-6 κB sites and reduced it at the IL-10 site (Fig. 4D–F, grey bars); p50 and p65 were both enriched at the TNF-α locus, whereas the IL-6 locus lost p50 but gained p65 (Fig. 4G,H,J,K, grey bars), consistent with p50 negatively regulating TNF-α but not IL-6 (*31*).

The central result here concerned the TNF-α κB site after strong TLR4 history, where TLR4 history and IL-10 signaling produced opposing, coordinated changes in factor occupancy. Strong priming drove BCL-3 accumulation that was maintained upon restimulation (Fig. 4D), while p65, recruited by acute stimulation, was instead reduced by prolonged strong stimulation and remained suppressed upon restimulation (Fig. 4J). Blocking the IL-10 receptor reversed both: BCL-3 occupancy fell by roughly two orders of magnitude and p65 occupancy was restored (Fig. 4D,J). Because cell-wide and nuclear BCL-3 levels changed only modestly under IL-10R blockade (Fig. S6D), the loss of promoter-bound BCL-3 likely reflects an IL-10-dependent change in its chromatin-binding competence rather than in its abundance. p50 was likewise enriched at the TNF-α κB site after prolonged stimulation but fell to basal levels upon acute restimulation, and, in the absence of IL-10 signaling, restimulation instead increased p50 alongside the IL-10-dependent gain of p65 (Fig. 4G,J). By contrast, the IL-6 and IL-10 loci showed more modest, largely IL-10-independent changes. The IL-6 κB site exhibited only subtle variation in p50 and p65 occupancy (Fig. 4H,K). At the IL-10 κB site, strong stimulation recruited p50, consistent with its role in promoting IL-10 expression. But restimulation did not further increase it. p50 enrichment was abolished by IL-10R blockade even though IL-10 production itself rose, indicating an interrupted autocrine feedback loop (Fig. 4I). BCL-3 enrichment at the IL-10 site was only slightly greater with intact than with blocked IL-10 signaling, suggesting an additional IL-10/STAT3-independent contribution to its recruitment there.

### History-gated IL-10 licensing enables trajectory-aware control of infection in a computational model

To explore and illustrate the possible functional consequences of the IL-10 licensing logic, we implemented an ordinary differential equation (ODE) model of a macrophage population responding to a pathogen challenge over a simulated time course of 10 days. Depending on the inflammatory response, pathogen levels may plateau after an initial rise and then get cleared or may escape (full description in Supplementary Methods Text S1).

TLR4 signaling has been explored in modeling work that focused on details of intracellular signaling (*50–52*) or more complex innate responses regulating, for instance, wound healing (*53*). Interestingly, among those prior molecular-level modeling studies was a qualitative assessment of pre-stimulation dependent adaptation based on negative feedback stemming from A20 production (*52*).

Compared to those studies, the model we used is rather simple as it is limited to exploring the different trajectories an infection may take depending on the type of phenomenological anti-inflammatory regulation the responding macrophages experience. The model assumes that macrophages are recruited in proportion to inflammation and, reflecting the experimental findings presented here, retain a quantitative memory of prior stimulation that establishes graded activation set points governing subsequent responsiveness. They re-activate only when the current stimulus matches or exceeds their remembered set-point, corresponding to the match-or-exceed behavior reported in Fig. 1. IL-10 is produced in proportion to the cells’ TLR4 exposure. To assess the effects of IL-10 licensing, we generated three variants of this model that, importantly, differ only in how IL-10 acts on the cellular inflammatory response: under history-gated licensing (variant 1), only cells that have committed to a strong TLR4 response are susceptible to IL-10. This variant implements history-gated IL-10 licensing on the basis of single-cell experimental data on the dependence of TNF-α and IL-10 production on TLR4 stimulation history (Figs. 2H, I, Fig. S2E, Fig. S7 and Supplementary Text 1 model notes). Accordingly, and in an interesting coincidence reported by our data, cells start producing IL-10 as soon as they can sense its regulatory effects. Under the concentration-only variant (variant 2), every cell is suppressed in proportion to the ambient IL-10 level; and under the no-regulation variant (variant 3), IL-10 has no effect.

The three model variants produce markedly different outcomes (Fig. 5A), and these differences are robust under parameter variation (Fig. S11 in Text S1). Under history-gated licensing (variant 1), naive cells respond to the rising pathogen and clear it within the first day (Fig. 5A, left panel; the teal curve is hidden beneath the red no-regulation curve, which clears the pathogen equally well). Inflammation and tissue damage rise transiently and then declined as the pathogen is cleared and committed cells are restrained by IL-10, settling at a low residual plateau (Fig. 5A, center and right panels). This incomplete resolution reflects the absence of an active resolution program in the base model: with no pro-resolving phenotype to clear accumulated damage, a low-grade damage-associated (DAMP) signal sustains residual inflammation. Under IL-10 concentration-only suppression (variant 2), ambient IL-10 mutes newly recruited cells before they can respond, leading to pathogen escape to its carrying capacity (Fig. 5A, left panel, orange curve) and sustained, pathogen-driven injury (Fig. 5A, right panel, orange curve). The tissue damage in this variant therefore stems from the pathogen load rather than from inflammation, which is suppressed. In the no-regulation variant (variant 3), the pathogen is cleared, but inflammation and tissue damage fail to resolve, locking the tissue into a chronic, self-sustaining inflammatory state (Fig. 5A, red curves). Note that, in the failure modes, comparable injury is caused by opposite mechanisms: unchecked pathogen in the concentration-only variant versus unresolved immunopathology in the no-regulation variant. Only history-gated licensing both controlled the pathogen and limited tissue damage. The reason is that newly recruited naive cells are not yet licensed and so remain free to respond to a rising challenge even in the presence of ambient IL-10, whereas IL-10 restrains only cells that have already committed to a strong response. The population therefore tracks the direction of the pathogen trajectory rather than the absolute IL-10 level, recapitulating the trajectory-aware behavior implied by our data (Fig. 1).

**Fig. 5:**
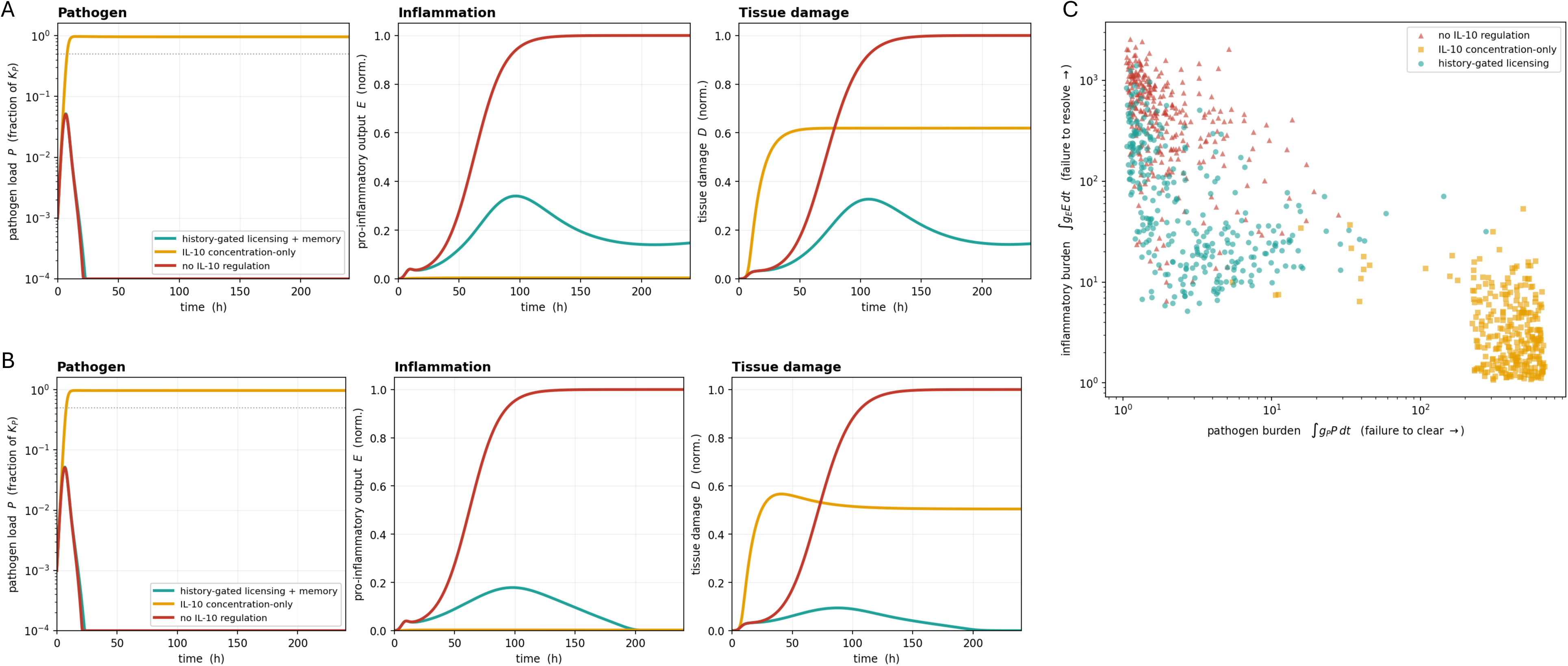
History-gated IL-10 licensing enables trajectory-aware control of infection in a computational model. (A) Base-model infection time courses for the three regulatory variants. Standard parameter set; history-gated licensing + memory (teal), IL-10 concentration-only (orange), no IL-10 regulation (red). Left panel: pathogen load (fraction of carrying capacity, log axis): concentration-only pre-mutes cells and the pathogen escapes to carrying capacity, whereas licensing+memory and no-regulation peak near 5% and clear it within ≈25 h. Center panel: pro-inflammatory output (normalized to its cross-variant maximum): absent in the escaped concentration-only case, transient-then-resolving under licensing+memory, sustained under no-regulation. Right panel: tissue damage (normalized): no-regulation locks into a non-resolving sterile-inflammation state, concentration-only incurs pathogen-driven injury, licensing+memory peaks then partially resolves. **(B) Active resolution by M1→M2 polarization.** Time courses with the resolution arm at the standard parameter set; colors as in (**A**). Left panel: pathogen (log axis); Center panel: inflammation; Right panel: tissue damage (center and right normalized to their cross-variant maxima). IL-10 now performs three coupled roles — acute suppression of licensed cells, the match-or-exceed memory gate, and driving the most-adapted cells into the M2-like state. Under licensing+memory the response resolves fully (final D → 0); no-regulation makes no M2 (IL-10R inert) and stays chronically inflamed; concentration-only again leads to pathogen escape. **(C) Two-objective decomposition of the parameter sweep.** Each point is one Latin-hypercube draw (resolution model with M2-like state), plotted by its pathogen burden (horizontal axis; rightward = failure to clear pathogen) against its inflammatory burden (vertical axis; upward = failure to resolve inflammation). The two alternatives to the history-gated licensing fail on opposite axes: concentration-only (orange) toward high pathogen burden, no-regulation (red) toward high inflammatory burden. History-gated licensing (teal) is the only variant that strongly populates the low-burden corner, clearing and resolving together.

In an extension of this model, and based on the gene-expression characteristics in cluster C3 of Fig. 2E, we further allowed the most strongly adapted, IL-10-suppressed macrophages to polarize toward a pro-resolving (M2-like) state that would eventually withdraw from inflammatory output and promote clearance of damaged tissue, reflecting the established coupling between IL-10 signaling and resolution programs (*54*). With this extension, the history-gated licensing variant fully clears the pathogen and resolves the inflammation, whereas the concentration-only variant still produces persistent tissue damage through its failure to clear the pathogen, and the no-regulation variant remains locked in uncontrolled inflammation (Fig. 5B).

While the model implements history-gated IL-10 licensing on the basis of single-cell experimental data, its kinetic parameters were not fitted to particular experiments. To assess the robustness of the licensing variant’s apparent advantage over the two alternatives, we performed parameter sweeps for which the parameter sets were drawn by Latin-hypercube sampling (*55*) to systematically cover the parameter space. Across the sampled space, the two alternatives to licensing failed along opposite axes: the concentration-only variant toward high pathogen burden and the no-regulation variant toward high inflammatory burden. Only the licensing variant consistently reached the low-burden region in which both pathogen and inflammation are controlled (Fig. 5C; per-parameter sensitivities from a Partial Rank Correlation Coefficient analysis can be found in Fig. S11). This structural separation suggests that the TLR4 stimulation memory and IL-10 licensing we identified in vitro may contribute to balancing pathogen clearance and tissue damage during macrophage responses to infection.

## Discussion

Macrophages and other innate immune cells must scale their inflammatory output to pathogen burdens that may change over time: responding vigorously when a threat escalates while limiting collateral damage to host tissue when it does not. By exposing BMDM to systematically varied sequences of KLA concentrations, we found that this balance is achieved, in part, through a quantitative memory of prior TLR4 activation and through a stimulation-history-dependent control of the cells’ own susceptibility to IL-10. Our data confirm established features of TLR4 sensitization and tolerance and the involvement of IL-10, BCL-3, p50 and p65 (*6–16, 19, 20*), but they additionally reveal three previously unrecognized aspects of macrophage adaptation.

First, MAPK and NF-κB signaling did not merely become blunted after prior stimulation but retained a quantitative memory of it: secondary responses required a stimulus that matched (p65) or exceeded (Erk1/2 and p38) the strength of the primary challenge, and the same match-or-exceed behavior propagated to the secreted pro-inflammatory mediators TNF-α, IL-6 and CXCL-1. Prolonged KLA exposure caused dose-dependent TLR4 endocytosis that did not reverse within several hours, so a reduced surface receptor pool likely contributes to adaptation of the MyD88 branch. Receptor loss alone, however, cannot account for the memory: even after internalization of ∼90% of surface TLR4, the macrophage population still discriminated the strength of a secondary stimulus. A recent systematic profiling of single-cell NF-κB activation dynamics across 80 pairwise ligand combinations at varied primary doses showed that the dose and duration of a prior inflammatory stimulus are encoded in subsequent NF-κB signaling, with both tolerance and priming arising in ligand- and dose-dependent ways (*10*). That work, however, varied only the primary stimulus dose against a single secondary dose, primarily across different TLR ligands. Here, we report a related but distinct form of memory: within a single TLR pathway, the amplitude of a secondary response is set by a quantitative comparison between secondary and primary stimulus strengths, and this match-or-exceed behavior extends from NF-κB dynamics to MAPK activation, IκBα degradation, and downstream cytokine and chemokine output. That this memory also governed TNF-α responses to intact, heat-inactivated gram-negative bacteria (Fig. S8) suggests it operates under physiologically relevant conditions.

Second, and central to this study, the cells’ susceptibility to IL-10 was not constitutive but was itself licensed by TLR4 stimulation history. Blocking the IL-10 receptor abolished the adaptation and memory of TNF-α and IL-6 production, whereas it left p38 and p65 phosphorylation largely unaffected, indicating that IL-10 acts downstream of the receptor-proximal kinase cascades.

Importantly, the requirement for strong prior TLR4 stimulation was not simply a consequence of greater IL-10 production following strong stimulation: titrating exogenous IL-10 to concentrations up to 100-fold above those generated by the cells themselves failed to effectively suppress TNF-α in the absence of strong priming, and, transcriptome-wide, an exogenous IL-10 concentration sufficient to suppress IL-6 altered only a small subset of genes in low-dose-primed macrophages. Consistent with IL-10 as the driver of adaptation, the switch of inflammatory signaling from low-dose sensitization to high-dose adaptation coincided with the onset of substantial IL-10 production. Direct, STAT3-mediated suppression of IL-6 thus appears to be the exception rather than a template for IL-10’s action; for most targets, including TNF-α and BCL-3, IL-10 responsiveness must be enabled by strong TLR4 signaling. This gating may have escaped notice in earlier work because IL-10’s effects on macrophages have typically been assayed with uniformly high LPS or KLA doses (*31, 37, 38*). The licensing we observed was graded rather than all-or-none: weakly (low-dose) primed macrophages were markedly less responsive to IL-10 than strongly primed cells, not wholly refractory to it. We also note that every condition we examined involved TLR4 engagement, our data do, thus, not address what IL-10 does to the other functions of fully naive macrophages that have never encountered a TLR4 ligand; the licensing we describe concerns the IL-10 susceptibility of the inflammatory program in TLR4-experienced cells.

Speculating on the physiological relevance of this licensing requirement, we think that several scenarios are consistent with our data, though none is directly tested here. Tissue macrophages routinely occupy microenvironments containing ambient IL-10, contributed by regulatory T cells, neighboring myeloid cells, and, at mucosal sites, steady-state epithelial microbiota-driven tone (*32, 33*). In such an environment, a constitutively IL-10-responsive cell would risk being silenced by this background before sensing a pathogen of its own. A second, complementary, pressure is pathogen-driven IL-10 evasion: multiple intracellular bacteria and protozoa induce host IL-10 (*34*), and several viruses encode IL-10 mimics that engage the host receptor (*35, 36*); generic IL-10 suppressibility would let such strategies silence the initial host response on encounter, whereas licensing buys the cell a response window before IL-10 can suppress it. The same gate may also help macrophage populations integrate divergent local trajectories: a resolving inflammatory focus adjacent to an expanding one, or a secondary infection superimposed on a primary one. In this setting, the rising arm could be handled by the quantitative memory described above and the falling arm by IL-10, but only once the cell has committed to a TLR response.

The most important role of IL-10 licensing may, however, be to allow for continuous evaluation of pathogen levels during an ongoing immune response. By tying a cell’s IL-10 susceptibility to its own TLR4 history, licensing lets each newly recruited macrophage respond to the pathogen levels it actually encounters, rather than being governed by cells that were activated earlier under potentially very different conditions.

What these scenarios share is the logic of an AND-gate that decouples IL-10 detection from IL-10 action until sufficient TLR engagement has occurred. Such a gate is potentially provided by the IL-10 and TLR4 stimulation history-dependent NFκB-mediated association of BCL-3 with the chromatin.

This chromatin-level mechanism for IL-10 licensing is the third major finding discussed here. An important detail of this finding is that neither the mRNA levels nor the nuclear accumulation of BCL-3, p50 or p65 predicted whether a target gene such as TNF-α would be suppressed, arguing against a simple mass-action competition among these factors. Chromatin immunoprecipitation instead revealed coordinated, IL-10-dependent changes in factor occupancy at the κB site of the

TNF-α gene: after strong TLR4 history, intact IL-10 signaling promoted recruitment of BCL-3 and a reciprocal loss of p65, and blocking the IL-10 receptor reversed both – reducing promoter-associated BCL-3 by roughly two orders of magnitude while cell-wide BCL-3 abundance changed only modestly. IL-10 therefore appears to condition the p50-mediated chromatin-association competence of BCL-3 rather than its abundance, providing a candidate molecular substrate for the history-dependent licensing of IL-10 responsiveness.

An important open question is whether the strength of TLR4 stimulation further sets chromatin accessibility at these loci. Recent work has shown that an initial inflammatory stimulus reshapes the global chromatin accessibility landscape in macrophages in a dose-dependent manner (*10*), and a TLR4-dose-dependent priming of accessibility specifically at κB sites of inflammatory genes would mechanistically unify the KLA concentration history with IL-10 signaling within a single regulatory step.

At the single-cell level, the history dependence of TNF-α and IL-10 was encoded differently. Strong stimulation recruited essentially the entire population into Tnf expression regardless of priming, yet the per-cell abundance of Tnf transcripts varied with stimulation history – weak-then-strong stimulation generating more high-expressing cells than dual strong stimulation, and closely resembling dual strong stimulation with the IL-10 receptor blocked. For IL10 mRNA, our data showed the converse: per-cell expression was largely invariant across protocols, so that total IL-10 output was set by the fraction of producing cells. These observations reconcile the apparently paradoxical secretion data and are consistent with population-level “quorum licensing” models of macrophage activation (*9, 47*).

Taken together, our findings indicate that macrophages register the history of a TLR4 challenge across several layers: surface receptor availability, the activation thresholds of MAPK and NF-κB signaling, and the chromatin association of BCL-3. Central to this adaptation is the cells’ adjustable, rather than fixed, responsiveness to anti-inflammatory IL-10, which, as discussed above, may enhance the robustness of macrophage responses when faced with complex antigen-and cytokine environments.

Framing tolerance and sensitization in these quantitative terms suggests that macrophages do not simply switch between “responsive” and “hypo-responsive” states but continuously gauge whether a pathogenic stimulus is escalating or receding and tune both their inflammatory output and their susceptibility to IL-10 accordingly. The mechanisms described here were defined in vitro using BMDM and the defined TLR4 ligand KLA. Their corroboration with intact bacteria notwithstanding, it will be important to test whether the same history-dependent licensing of IL-10 operates in tissue macrophages in vivo, for example, by asking whether IL-10-rich mucosal niches selectively spare TLR4-unprimed cells from IL-10-mediated suppression.

Our experiments identify cell-intrinsic mechanisms shaping how macrophages may sense evolving pathogen loads, combining quantitative memory of TLR4 stimulation with history-gated susceptibility to IL-10. But they do not illustrate how this logic matters at the scale of an infected tissue. Our computational model is a step toward addressing this.

When IL-10 acts only on cells that have committed to a strong TLR4 response, the macrophage population tracks the direction of the pathogenic challenge, mounting a response against a rising load and resolving it as the load falls. When IL-10 instead acts on every cell in proportion to its concentration, newly recruited cells are silenced before they can read their own input. As a consequence, the information whether the pathogen load is rising or falling is lost and the pathogen may escape due to inappropriate suppression of inflammatory output. Removing IL-10 regulation altogether avoids that failure but yields unresolved, chronic inflammation. History-gated licensing escapes this trade-off rather than balancing it: by restricting IL-10’s reach to already-committed cells, it reaches the regime in which the pathogen is controlled and immunopathology is limited, satisfying within a single mechanism both demands, pathogen resistance and tolerance, that the two alternatives can only trade against each other.

The model is conceptual rather than predictive: it is dimensionless aside from a mapping of its time axis to hours, and, even though it reproduces the typical time course of an innate response, is not fitted to in-vivo infection kinetics. Its qualitative conclusions are nonetheless robust to wide parameter variation (Fig. 5C and S10) and are strengthened by the inclusion of an active resolution step (Fig. 5B). The framework also illustrates the cost of indiscriminate suppression: because pathogen escape forecloses the early clearance window, no downstream resolution program can compensate for it. This offers a functional rationale for why susceptibility to IL-10 is gated by TLR4 history rather than imposed uniformly. This framework predicts that ambient IL-10, including IL-10 homologs encoded by certain pathogens, would be considerably less effective as an immune-evasion strategy in a history-gated licensing system than in one governed solely by cytokine concentration

## Materials and Methods

### Materials

Kdo2-Lipid A (KLA) was purchased from Avanti Polar Lipids. Mek inhibitor U0126 (V1121) was obtained from Promega and re-constituted in DMSO (Sigma-Aldrich, D8418). The Alexa Fluor™ 647-conjugated threonine 202- and tyrosine 204-phosphorylated Erk1/2 (13148), Pacific Blue™- conjugated IκBα (13656), Alexa Fluor™ 647-conjugated serine 536-phosphorylated NF-κB p65 (4887) and PE-conjugated NF-κB1 p105/p50 (24961) antibodies were obtained from Cell Signaling Technology. PE-CF594-conjugated threonine 180- and tyrosine 182-phosphorylated p38-MAPK (563569) and PE-conjugated tyrosine 705-phosphorylated STAT3 (612569) antibodies were purchased from BD Biosciences. FITC-conjugated BCL-3 (LS-C62564) antibody was bought from LSBio. The PE-conjugated antibody for murine TLR-4 (145404) was purchased from BioLegend. anti-TNFR1 (Armenian hamster IgG, 16-1202-85), its isotype control (Armenian hamster IgG, 16-4888-85), anti-IL-10R (rat IgG1κ, 16-2101-85), its isotype control (rat IgG1κ, 14-4301-85), and anti-IFNAR1 (mouse IgG1κ, 16-5945-85), its isotype control (mouse IgG1κ, 14-4714-85) were from Invitrogen. Dulbecco’s modified Eagle’s medium (DMEM) was from Gibco Life Technologies. Heat-inactivated fetal bovine serum (FBS) was bought from GeminiBio.

### Preparation of Heat-inactivated Bacteria

Bacterial strain used in this study included main model *E.coli* strain K12 from ATCC (K12 MG1655). Bacteria were struck out on a Luria-Bertani (LB) agar plate and grown overnight (ON) to isolate single colonies. Single colony was frozen in a 25% glycerol stock. Bacterial culture was grown from glycerol stock with shaking at 220 rpm at 37 °C ON to saturation phase in LB broth media (KD Medical, catalog #: BLF-7030), then adjusted based on OD600 to the indicated MOI, washed 3x with PBS to clear any debris left in the media from the ON, and finally resuspended in PBS. Bacteria were heat-inactivated at 100 °C for 30 min prior to use.

### Cells and Cell Culture

Mice were maintained in specific-pathogen-free conditions at an American Association for Laboratory Animal Care-accredited animal facility at the National Institute of Allergy and Infectious Diseases (NIAID, NIH) and were used under study protocol LISB-4E approved by the NIAID Animal Care and Use Committee (National Institutes of Health, NIH).

Bone marrow cells were harvested from femurs and tibias of C57BL/6J (JAX664) mice and bone marrow progenitors were plated on non-tissue treated dishes and differentiated into BMDM during a 6-day culture in complete Dulbecco’s modified Eagle’s medium (DMEM) containing 5% FBS and supplemented with 60 ng/ml recombinant murine M-CSF (Stemcell Technologies) in a water saturated atmosphere at 37°C. For experiments, 2.5×10^6^ BMDM were seeded on non-tissue treated wells of 48 well plates in 5% FBS containing DMEM. On day of experiment, medium was replaced with fresh 5% FBS containing DMEM 30 min before experimental treatment.

### Flow cytometric analyses

FACS analysis of cell surface markers was conducted by placing cells on ice, replacing the medium with 5 mM EDTA/PBS buffer, and transferring them into a 96 well plate. Cells were blocked with 2% FBS and 0.2% goat serum containing HBSS including 1:1000 Mouse BD Fc Block™ (Clone 2.4G2, BDBiosciences) and stained with cell surface marker detecting fluorophore-conjugated antibodies. FACS analysis of intracellular proteins was performed by fixing cells with 2.5% PFA for 10 min at room temperature, permeabilizing in 90% methanol for 30 min at −30°C, and blocking with 2% FBS and 0.2% goat serum containing HBSS including 1:1000 Mouse BD Fc Block™ (Clone 2.4G2, BDBiosciences). Intracellular proteins were stained with fluorophore-conjugated antibodies overnight at 4°C. Samples were analyzed using a LSRII flow cytometer (BDBiosciences). The percent maximum value was calculated by subtraction of the baseline MFI value and division of the MFI of each sample by the maximal MFI for each protein species measured within one experiment, multiplied by 100.

### Quantitative real-time PCR

Total RNA was isolated using QiaShredder columns and the RNeasy Mini Kit (Qiagen) according to manufacturer’s instructions. RNA (100 ng) was reverse transcribed into cDNA with iScript cDNA synthesis kit (Bio-RAD) according to manufacturer’s instructions. Gene expression of murine BCL-3, cFos, cJun, CCL-2, CXCL-1. CXCL-10, IL-6, IL-10, IFN-β, Junb, TNF-α, SOCS3 and SDHA was measured with primers for murine CXCL-1 (fw: 5′- GCT TGA AGG TGT TGC CCT CAG -3′, rev: 5′- AAG CCT CGC GAC CAT TCT TG -3′), murine CXCL-10 (fw: 5′- GCC GTC ATT TTC TGC CTC AT -3′, rev: 5′- GCT TCC CTA TGG CCC TCA TT -3′), murine CCL-2 (fw: 5′- TTA AAA ACC TGG ATC GGA ACC AA -3′, rev: 5′- GCA TTA GCT TCA GAT TTA CGG GT -3′), murine IL-6 (fw: 5′-CTC TGC AAG AGA CTT CCA TCC AGT -3′, rev: 5′-GAA GTA GGG AAG GCC GTG G -3′), murine IL-10 (fw: 5′- AAG GCA GTG GAG CAG GTG AA -3′, rev: 5′- CCA GCA GAC TCA ATA CAC AC -3′), murine IFN-β (fw: 5′- AAG AGT TAC ACT GCC TTT GCC ATC -3′, rev: 5′- CAC TGT CTG CTG GTG GAG TTC ATC -3′), murine TNF-α (fw: 5′- CAT CTT CTC AAA ATT CGA GTG ACA A -3′, rev: 5′- TGG GAG TAG ACA AGG TAC AAC CC -3’), murine BCL-3 (fw: 5′- GGA GCC GCG AAG TAG ACG T - 3′, rev: 5′- TGT GGT GAT GAC AGC CAG GT -3′), murine SOCS3 (fw: 5′- GCT CCA AAA GCG AGT ACC AGC -3′, rev: 5′- AGT AGA ATC CGC TCT CCT GCA G -3′), murine JunB (fw: 5’-ATG TGC ACG AAA ATG GAA CA-3’, rev: 5’-CCT GAC CCG AAA AGT AGC TG-3’), murine cFos (fw: 5’-CGA AGG GAA CGG AAT AAG ATG-3’, rev: 5’-GCT GCC AAA ATA AAC TCC AG-3’), murine cJun (fw: 5’-AAA ACC TTG AAA GCG CAA AA-3’, rev: 5’-CGC AAC CAG TCA AGT TCT CA-3’),and murine SDHA (fw: 5’- TGG GGA GTG CCG TGG TGT CA - 3’, rev: 5’- GTG CCG TCC CCT GTG CTG GT -3’). Real-time PCR was performed using Fast SYBR™ Green Master Mix (ThermoFisher Scientific, USA) according to manufacturer’s instructions in MicroAmp™ Fast Optical 96-Well Reaction Plates ThermoFisher Scientific, 4346906) with a QuantStudio™ 6 Flex Real-Time PCR System (ThermoFisher Scientific, 4485691). Quantification of gene expression was calculated as described by Pfaffl et al. (*56*).

### Immunocytochemistry and widefield fluorescence microscopy

For widefield fluorescence microscopy, 1×10^5^ BMDM cells were seeded on Falcon 96 well Flat Bottom TC-treated Imaging Microplate (Corning, 353219) and cultivated for 24h before experimental treatment. After treatment, cells were fixed with 2.5% PFA for 10 min at room temperature. Cells were then permeabilized in 70% ethanol for 30 min. After permeabilization, cells were blocked with 2% FBS and 0.2% goat serum containing HBSS including 1:1000 Mouse BD Fc Block™ (Clone 2.4G2, BDBiosciences). For detection of endogenous p65, BCL-3 and p50, cells were stained with antibodies specific for p65, BCL-3 and p50 overnight at 4°C. Additionally, DAPI (BioLegend) was added for nuclear staining. Samples were imaged using a CellInsight CX7 Pro HCS Platform (ThermoFisher Scientific) equipped with an 40x objective lens. Quantification of the nuclear localization of Alexa Fluor-647-labeled p65, FITC-labeled BCL-3 and PE-labeled p50 were performed by using CellProfiler™ for determining the total intensity of fluorophores inside the nucleus, and the relative ratio of intensity of the fluorophores in the nucleus and in the cytoplasm. Nuclear to cytoplasmic ratio was calculated as the nuclear localized fluorophore intensity divided by the intensity of the fluorophore in the cytoplasm.

### Cytokine release quantification

For assessing cyto- and chemokine release, supernatant of treated BMDM was taken after putting cells on ice. Supernatant was frozen at -80°C for long-term preservation. After thawing, supernatant was prepared and analyzed with the LegendPlex™ Multiplex Assay Kits Mouse Proinflammatory Chemokine Panel (740451) and Mouse Inflammation Panel (740446) according to manufacturer’s instructions by using a LSRII flow cytometer (BDBiosciences). Raw data was further processed with LegendPlex™ Desktop software.

### Chromatin immunoprecipitation (ChIP) assay

Samples for ChIP assays and sequencing from in vitro cultured BMDM have been prepared according to Rousselet (*57*). After fixation of adherent cells on a 10 cm dish in 1% paraformaldehyde and adding glycine to a final concentration of 125 mM, cells were washed with PBS containing phosphatase and protease inhibitor cocktail Halt™ (1:500) (ThermoFisher Scientific, 78440), transferred in a 15 ml tube and centrifuged for 8 min at 300x g and 4 °C. Pellet was resuspended in SDS-containing lysis buffer and lysates have been passed through a 27Gx1/2 needle to ensure separation of nuclei. Lysates were sonicated with a BioRuptor UCD-300 (Diagenode) for 13 cycles of 30 s on/30 s off, respectively. This was repeated 3 times. To check for the extent of chromatin shearing, 10% of the sample were transferred in a new tube and treated with Proteinase K and RNase and were reverse crosslinked. DNA was then extracted with phenol/chloroform/isoamylalcohol and precipitated with ethanol. DNA was size fractionated via gel electrophoresis on a 1% agarose gel. Successful chromatin fragmentation resulted in a smear of DNA fragments ranging from 100-600 bp. Fragmented chromatin has been pelleted and resuspended in ChIP dilution buffer (see (*57*)). Samples were pre-cleared with Pierce™ ChIP-grade Protein A/G conjugated magnetic beads (ThermoFisher Scientific, 26162). Beads were removed using a magnetic tube rack and supernatants containing the pre-cleared chromatin were splitted into immunoprecipitation (IP) and input samples. Supernatants of each sample that were used for IP, have been incubated with 10 μg of either anti-BCL3 (SantaCruz, sc-32741) or it’s corresponding isotype control antibody (Armenian Hamster IgG, ThermoFisher Scientific, 16-4888-85) overnight at 4 °C. p50 and p65 IP have been performed with anti-p50 (Santa Cruz, sc-8414X) or anti-p65 (Santa Cruz, sc-8008X) antibodies or their respective isotype control antibody (Mouse IgG1 kappa, ThermoFisher Scientific, 14-4714-85). The next day, in 1.5% fish skin gelatin (Sigma-Aldrich, G7041) and glycogen (Sigma-Aldrich, G8751) pre-blocked Pierce™ ChIP-grade Protein A/G conjugated magnetic beads (ThermoFisher Scientific, 26162) were added to IP samples to bind chromatin/antibody complexes and incubated for 2 h at 4 °C. Tubes were placed in a magnetic tube rack, liquid was removed and beads were washed with a succession of buffers according to Rousselet (*57*). Chromatin/antibody complexes were eluted from the beads for 30 min. IP samples and Input samples were reverse crosslinked over night at 65°C in a rotary shaker. After adding Proteinase K and RNase, samples were incubated for 1 h at 37 °C. DNA was purified using Qiaquick® PCR Purification Kit (Qiagen, 28104).

For analysis using quantitative real-time PCR, IP DNA sample and input sample were used as templates. Real-time PCR analysis of DNA samples was performed according to procedure described in section ‘Quantitative real-time PCR’ using primer for known BCL3 specific κB-sites in the murine TNF-α gene (fw: 5’-CCA GGA GGG AGA ACA GAA ACT C-3’, rev: 5’- CAC AAG CAG GAA TGA GAA GAG G -3’) (*31*), murine IL-6 gene (fw: 5’- GAC ATG CTC AAG TGC TGA GTC AC -3’, rev: 5’- AGA TTG CAC AAT GTG ACG TCG -3’) (*58*) and murine IL-10 gene (fw: 5’- TAG AAG AGG GAG GAG GAG CC -3’, rev: 5’- TGT GGC TTT GGT AGT GC AAG -3’) (*59*). Data was analyzed and is given as ratio (fold enrichment) comparing the amount of the target sequence measured in the ChIP isolate to the amount measured in the isotype control isolate.

### Bulk RNA-seq

For bulk RNA sequencing, total RNA from BMDM was isolated using QiaShredder columns and the RNeasy Mini Kit (Qiagen) according to manufacturer’s instructions. Before eluting isolated RNA from RNeasy spin columns, bound RNA/DNA has been incubated with DNase I (RNase-Free DNase Set, Qiagen) to remove DNA from samples. After RNA elution, RNA quantity and integrity was assessed by the Bioanalyzer (Agilent). 100 ng of total RNA was used in conjunction with the TruSeq® Stranded Total RNA Library Prep kit (Illumina). The libraries quality was checked by the Bioanalyzer and quantitated by the Qubit (ThermoFisher Scientific). Equimolar quantities from each sample library were pooled and sequenced on a High throughput Next-Seq2000 instrument.

### RNA sequencing data analysis

Paired-end sequence files (.fastq) per sample were quality inspected using the FastQC tool 0.12.1 (https://www.bioinformatics.babraham.ac.uk/projects/fastqc/) Reference mapping of each fastq-file has been performed with the Rhisat2 package (https://bioconductor.org/packages/release/bioc/html/Rhisat2.html) in R using the mouse reference genome GRCm38 (https://cloud.biohpc.swmed.edu/index.php/s/grcm38/download).

Gene expression has been quantified after annotation of mapped genes from the saf file provided by https://bioinf.wehi.edu.au/Rsubread/annot/ and after counting reads to genes from the genomic alignment with the Rsubread package (https://bioconductor.org/packages/release/bioc/html/Rsubread.html) in R. Counts have been Z- score normalized. For differential gene expression (DEG) analysis, genes with less than 10 reads have been excluded. For genes not discarded, expression differences across sample classes were tested using the DESeq2 (https://bioconductor.org/packages/release/bioc/html/DESeq2.html) package in R.

### Single cell RNA sequencing

Single cell suspensions were collected into PBS+0.5% BSA. Single cell suspensions quality, number and viability were assessed with a dual fluorescence cell counter LUNA-FX7TM (Logos Biosystems). 3,000 cells were targeted from each sample cell suspension. The cells were washed twice with PBS+0.04% BSA and resuspended in about 500 cells per microliter.

10X Genomics’ Chromium instrument and Dual index Single-Cell 3′ Reagent kit (V3.1) were used to prepare the individually barcoded single-cell RNA-Seq libraries following the manufacturer’s protocol. Library qualities were assessed by the TapeStation-4200 traces (Agilent BioAnalyzer High Sensitivity Kit) and quantitated by the Qubit system (ThermoFisher Scientific). Sequencing was done on the Illumina NextSeq-2000 machine, using the P3 kit. Following sequencing, an average 50,000 reads per cell were generated. The bcl files were demultiplexed into a FASTQ, aligned to Mouse transcriptome mm10-2020-A and single-cell 3′ gene counting were performed by the standard 10X Genomics’ CellRanger mkfastq software (V8.0.0).

The single-cell QC and the 10X Genomics’ H5 output files, containing the gene counts per cell matrix were generated by 10X Genomics’ CellRanger Downstream analysis utilized a Python-based pipeline that primarily relied on Scanpy (v1.9.8) for single cell dataset processing. Briefly, H5 count matrices from all samples were simultaneously loaded in using Anndata, and duplicate indices across experiments were given unique names. Mitochondrial-derived genes were identified and used for standard Scanpy quality control checks. After confirming satisfactory quality of the experimental data, the count matrix was filtered to exclude cells that expressed less than 100 genes and genes which were expressed in fewer than 3 cells. This filtered matrix was then subject to normalization using the Scanpy normalize_total function, which normalizes the total counts per cell across the dataset. This normalized matrix was then scaled using a log+1 function, and these filtered, normalized, and scaled matrices were lastly further filtered to only include the 20,000 most highly variable genes. These final matrices were then used for all downstream analyses.

After this standard Scanpy pipeline, analysis was done using a custom script. UMAP plots were created using the Scanpy (v1.9.8) package, with figures generated using matplotlib. Expression node figures were created in Adobe Illustrator based on a simple binary determination of cellular gene expression. Counts of cells expressing each given gene were recorded, while simultaneously keeping track of cellular identity to determine single, double, triple, and quadruple positive cells for given marker genes.

### Standard statistical analyses

Statistical analyses were performed with R (version 4.3.0) and RStudio (version 2023.03.1) for MacOS. Data were tested for normal distribution with the Shapiro-Wilk test and homoscedasticity using Levene test. In case of non-normally distributed or heteroskedastic data, nonparametric tests such as Mann–Whitney-U (for single comparison) and Kruskal–Wallis (followed by post hoc Dunn–Bonferroni for multiple comparisons) were applied. For normally distributed data, parametric tests such as *t*-test (for single comparison) and ANOVA (followed by post hoc Tukey for multiple comparisons) were performed. The designations * p < .05, ** p < .01, *** p < .001 denote p-values for the measured differences. If no p-value is indicated, the comparison is considered non-significant. All experiments contained a minimum of three replicates (n = 3).

### Computational model of macrophage population responses

See Supplementary Text 1.

## Acknowledgements

We thank the members of the Laboratory of Immune System Biology (LISB), in particular Ronald Germain, for very helpful comments on this project and the manuscript. We also thank Dorian McGavern (Viral Immunology and Intravital Imaging Section, National Institute of Neurological Disorders and Stroke (NINDS)), Abdel G. Elkahloun and Bayu Sisay (Microarrays and Single-cell Genomics Core, National Human Genome Research Institute (NHGRI)) for providing valuable assistance and support in RNA-seq experiments.

## Author Contributions

Conceptualization: H.B., B.M. and M.M.-S.; Methodology: H.B., B.M., C.B., J.G., R.A.G., A.N.-L., I.D.C.F., M.M.-S.; Formal analysis: H.B., C.T.B., T.P., M.M-S.; Investigation: H.B., B.M.; Drafting original manuscript: H.B. and M.M-S.; Review and editing: all authors.

## AI usage statement

Claude 4.8 was used to support literature reviews and to suggest text passages that could be tightened. It was also used to generate Python versions of the simulations of the model and, in particular, to help with generating figures showing simulation results.

## Figure legends / specific methods

**Fig. S1:**
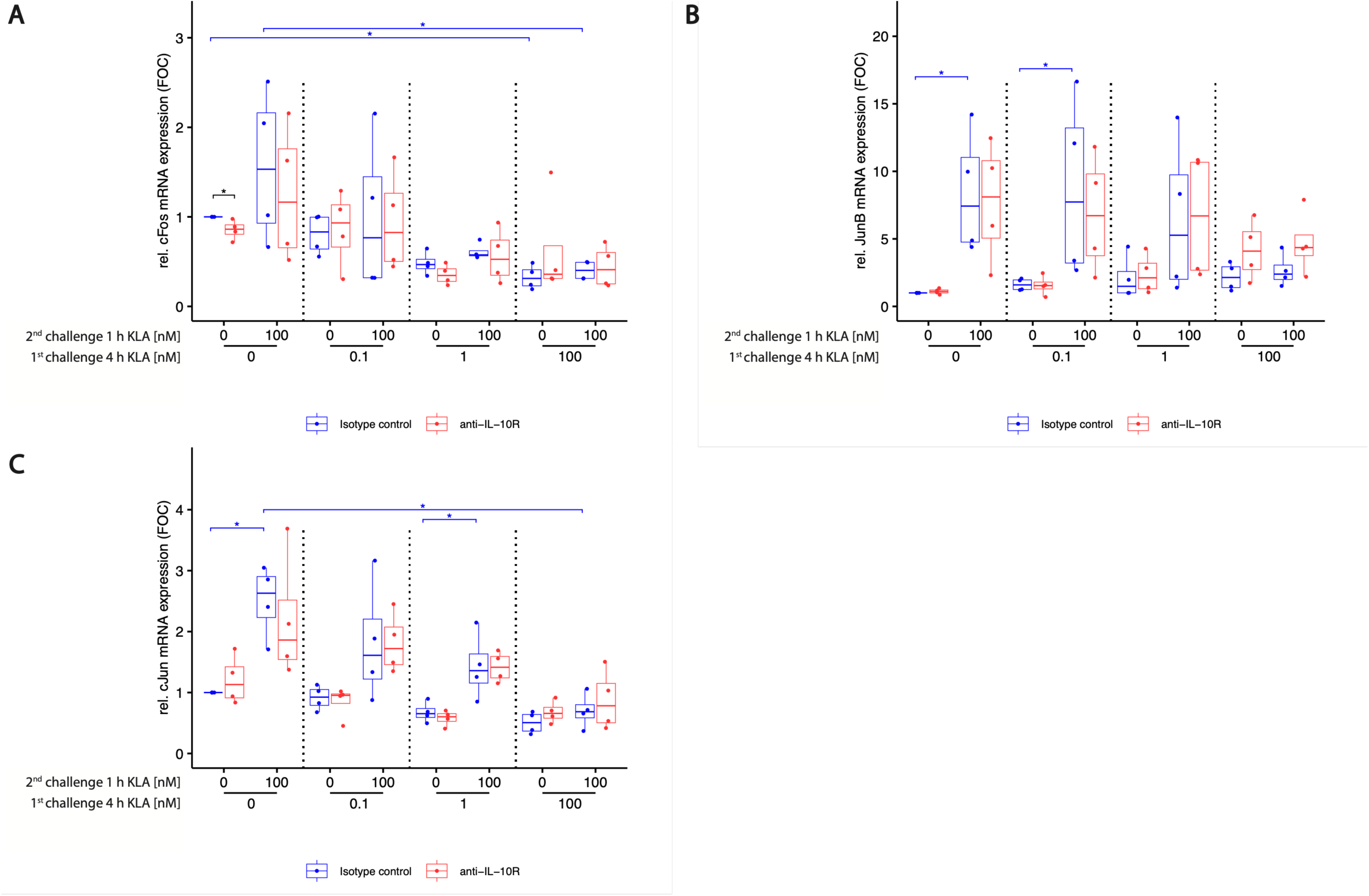
Blocking IL-10 does not affect AP-1 family mRNA expression in TLR4 ligand pre-exposed and re-stimulated macrophages. BMDMs were incubated with 5% FBS DMEM containing either IL-10R blocking antibody or its isotype control. During primary stimulation, cells were stimulated with 0, 0.1, 1 or 100 nM KLA. After 4 hrs, cells were washed, and medium was replaced with 5% FBS DMEM containing either IL-10R blocking antibody or its isotype control and cells were incubated for 1 hr. Subsequently, re-stimulation was performed using 0 or 100 nM KLA. After 1 hr, total RNA was isolated and subjected to qRT-PCR analysis to monitor (A) cFos, (B) JunB and (C) cJun mRNA expression. The expression of aforementioned mRNAs was normalized to SDHA mRNA expression. Relative expression of mRNA is given in fold of mRNA expression in naïve and unstimulated control cells (fold of control (FOC)) as described by Pfaffl et al. (*56*). Each replicate was performed with BMDMs from different mice and data sets include 4 replicates (n=4). Kruskal–Wallis test with post-hoc Dunn-Bonferroni comparisons: \**p* < .05, \*\**p* < .01, \*\*\**p* < .001, \*\*\*\**p* < .0001.

**Fig. S2:**
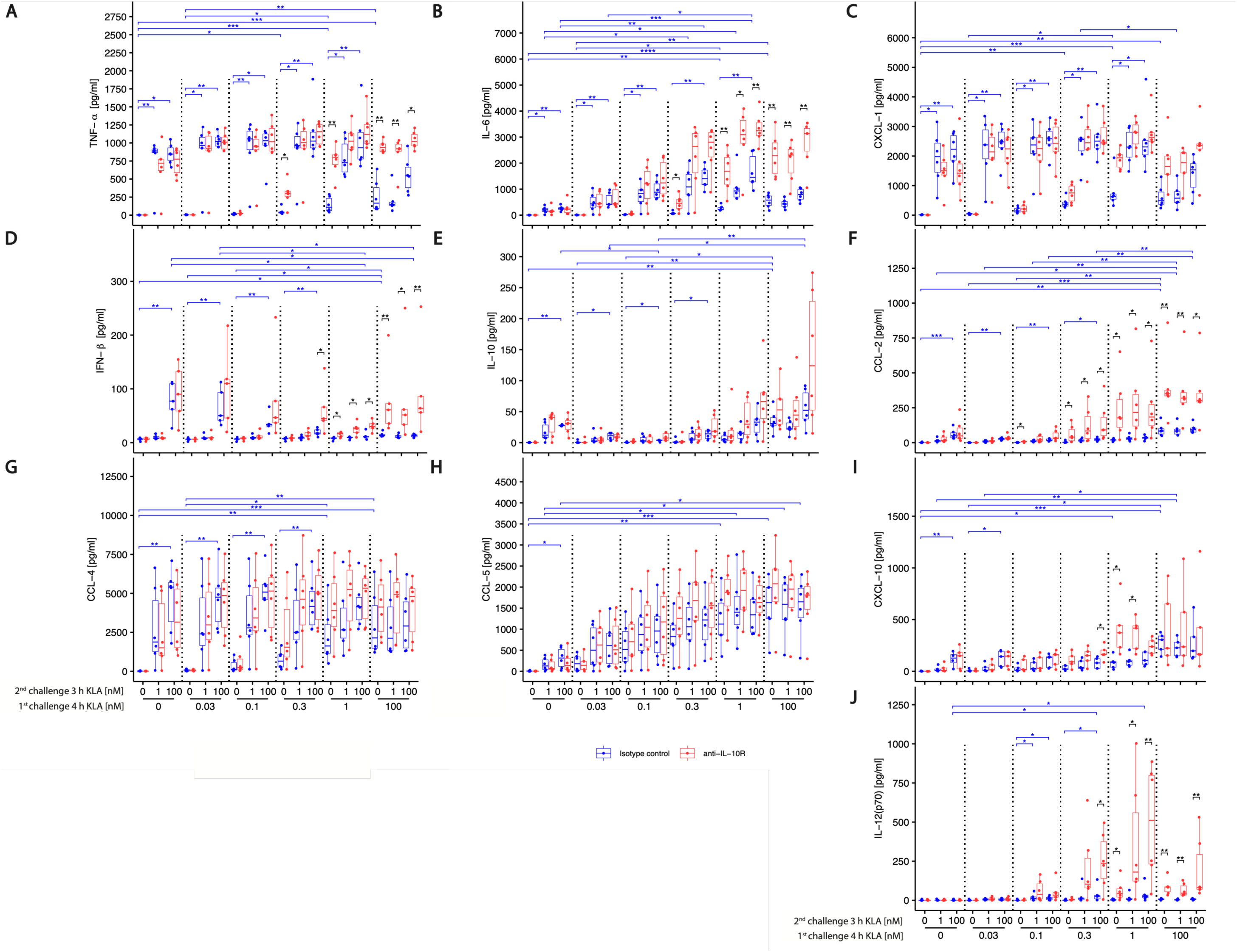
Low dose KLA priming dependent increase in restimulation induced cyto- and chemokine release is accompanied by reduced IL-10 expression. BMDMs were incubated with 5% FBS DMEM containing either IL-10R blocking antibody or its isotype control. During primary stimulation, cells were treated with 0, 0.03, 0.1, 0.3, 1, or 100 nM KLA. After 4 hrs, cells were washed, and medium was replaced with 5% FBS DMEM containing either IL-10R blocking antibody or its isotype control and cells were incubated for 1 hr. Subsequently, restimulation was performed using 0, 1 or 100 nM KLA. After 3 hrs, supernatants were collected, processed with the LegendPlex™ Multiplex Assay Kits and cyto- and chemokine levels for (A) TNF-α, (B) IL-6, (C) CXCL-1, (D) IFN-β, (E) IL-10, (F) CCL-2, (G) CCL-4, (H) CCL-5, (I) CXCL-10, and (J) IL-12p70 were determined by using a LSRII flow cytometer (BDBiosciences). Raw data was processed with LegendPlex™ Desktop software. Each replicate was performed with BMDMs from different mice and data sets include 6 replicates (n=6). Data was normalized to the maximal cytokine secretion (set as 100%). Kruskal–Wallis test with post-hoc Dunn-Bonferroni comparisons: \**p* < .05, \*\**p* < .01, \*\*\**p* < .001, \*\*\*\**p* < .0001.

**Fig. S3:**
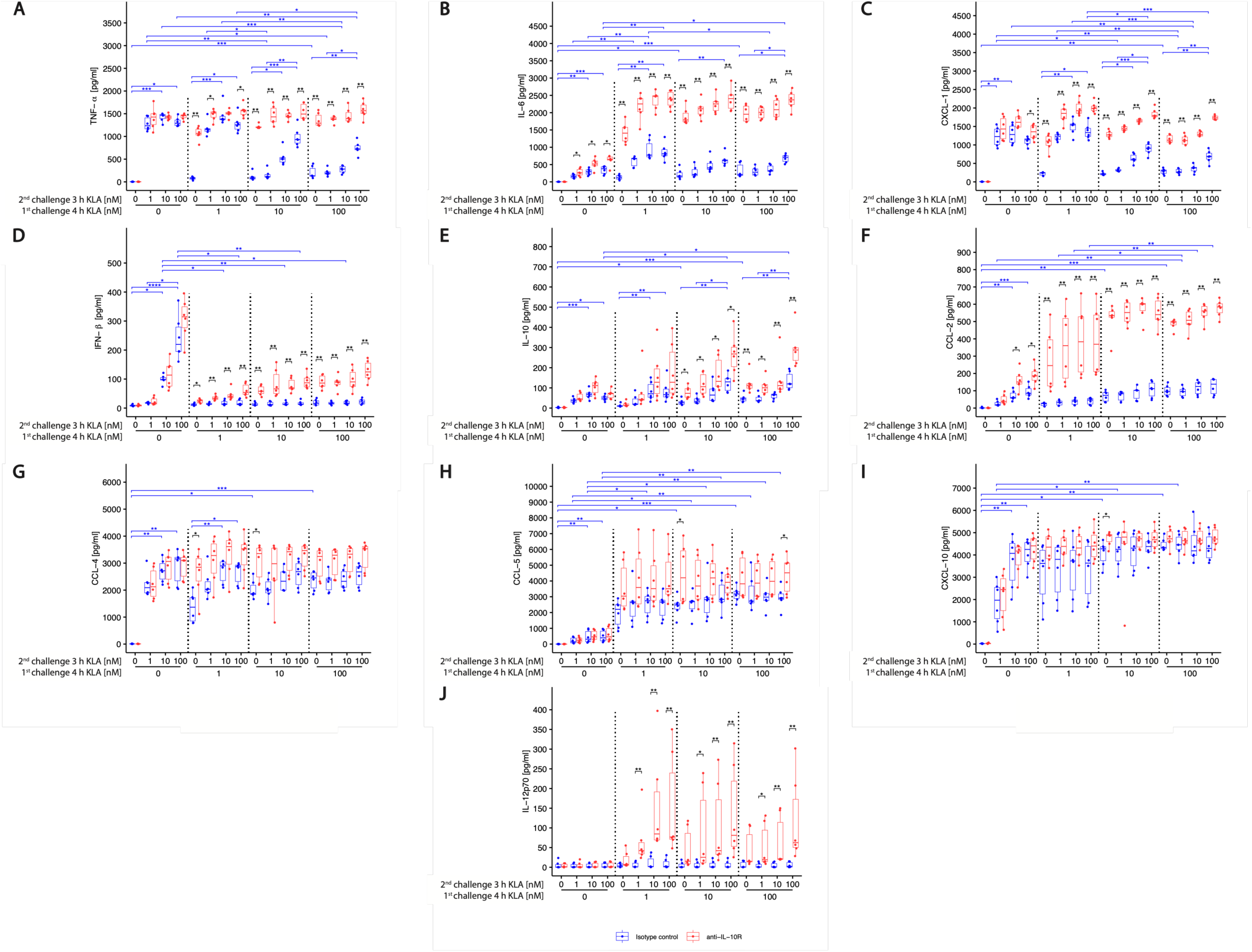
High dose KLA primary stimulation induces hyporesponsive behavior of cytokine and chemokine release in response to restimulation while blocking IL-10 reverses hyporesponsiveness. BMDMs were incubated with 5% FBS DMEM containing either IL-10R blocking antibody or its isotype control. During primary stimulation, cells were treated with 0, 1, 10 or 100 nM KLA. After 4 hrs, cells were washed, and medium was replaced with 5% FBS DMEM containing either IL-10R blocking antibody or its isotype control and cells were incubated for 1 hr. Subsequently, restimulation was performed using 0, 1, 10 or 100 nM KLA. After 3 hrs, supernatants were collected, processed with the LegendPlex™ Multiplex Assay Kits and cyto- and chemokine levels for (A) TNF-α, (B) IL-6, (C) CXCL-1, (D) IFN-β, (E) IL-10, (F) CCL-2, (G) CCL-4, (H) CCL-5, (I) CXCL-10, and (J) IL-12p70 were determined by using a LSRII flow cytometer (BDBiosciences). Raw data was processed with LegendPlex™ Desktop software. Each replicate was performed with BMDMs from different mice and data sets include at least 5 replicates (n=5). Kruskal–Wallis test with post-hoc Dunn-Bonferroni comparisons: \**p* < .05, \*\**p* < .01, \*\*\**p* < .001, \*\*\*\**p* < .0001.

**Fig. S4:**
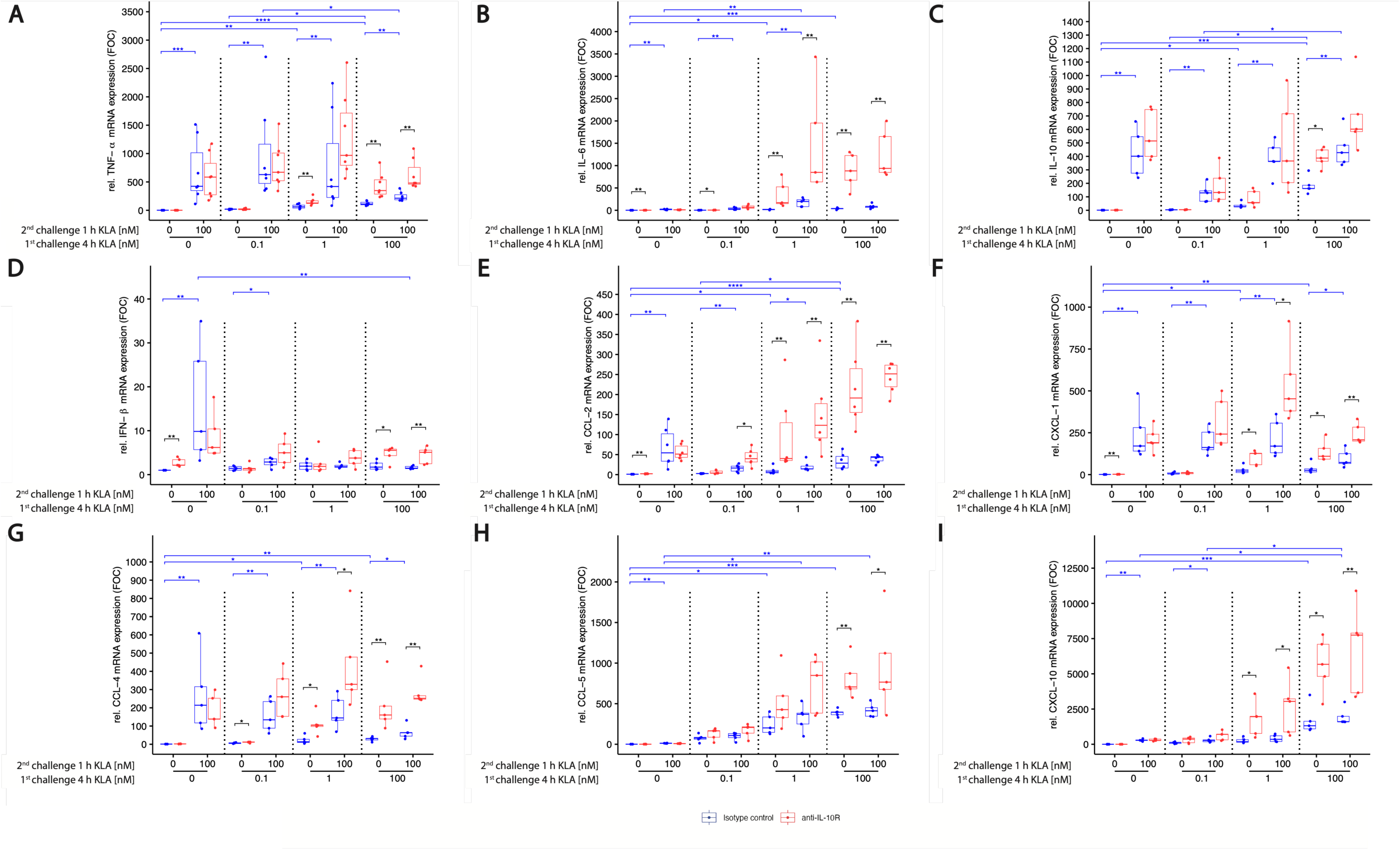
KLA concentrations during primary stimulations affect RNA production of cytokines and chemokines upon restimulation. Blocking IL-10 reverses hypo-responsiveness. BMDMs were incubated with 5% FBS DMEM containing either IL-10R blocking antibody or its isotype control. During primary stimulation, cells were treated with 0, 0.1, 1 or 100 nM KLA. After 4 hrs, cells were washed, and medium was replaced with 5% FBS DMEM containing either IL-10R blocking antibody or its isotype control and cells were incubated for 1 hr. Subsequently, re-stimulation was performed using 0 or 100 nM KLA. After 1 hr, total RNA was isolated and subjected to qRT-PCR analysis to monitor (A) TNF-α, (B) IL-6 (C) IL-10, (D) IFN-β, (E) CCL-2, (F) CXCL-1, (G) CCL-4, (H) CCL-5 and (I) CXCL-10 mRNA expression. The expression of aforementioned mRNAs was normalized to SDHA mRNA expression. Relative expression of mRNA is given in fold of mRNA expression in naïve and unstimulated control cells (fold of control (FOC)) as described by Pfaffl et al. (*56*). Each replicate was performed with BMDMs from different mice and data sets include at least 5 replicates (n=5). Kruskal–Wallis test with post-hoc Dunn-Bonferroni comparisons: \**p* < .05, \*\**p* < .01, \*\*\**p* < .001, \*\*\*\**p* < .0001.

**Fig. S5:**
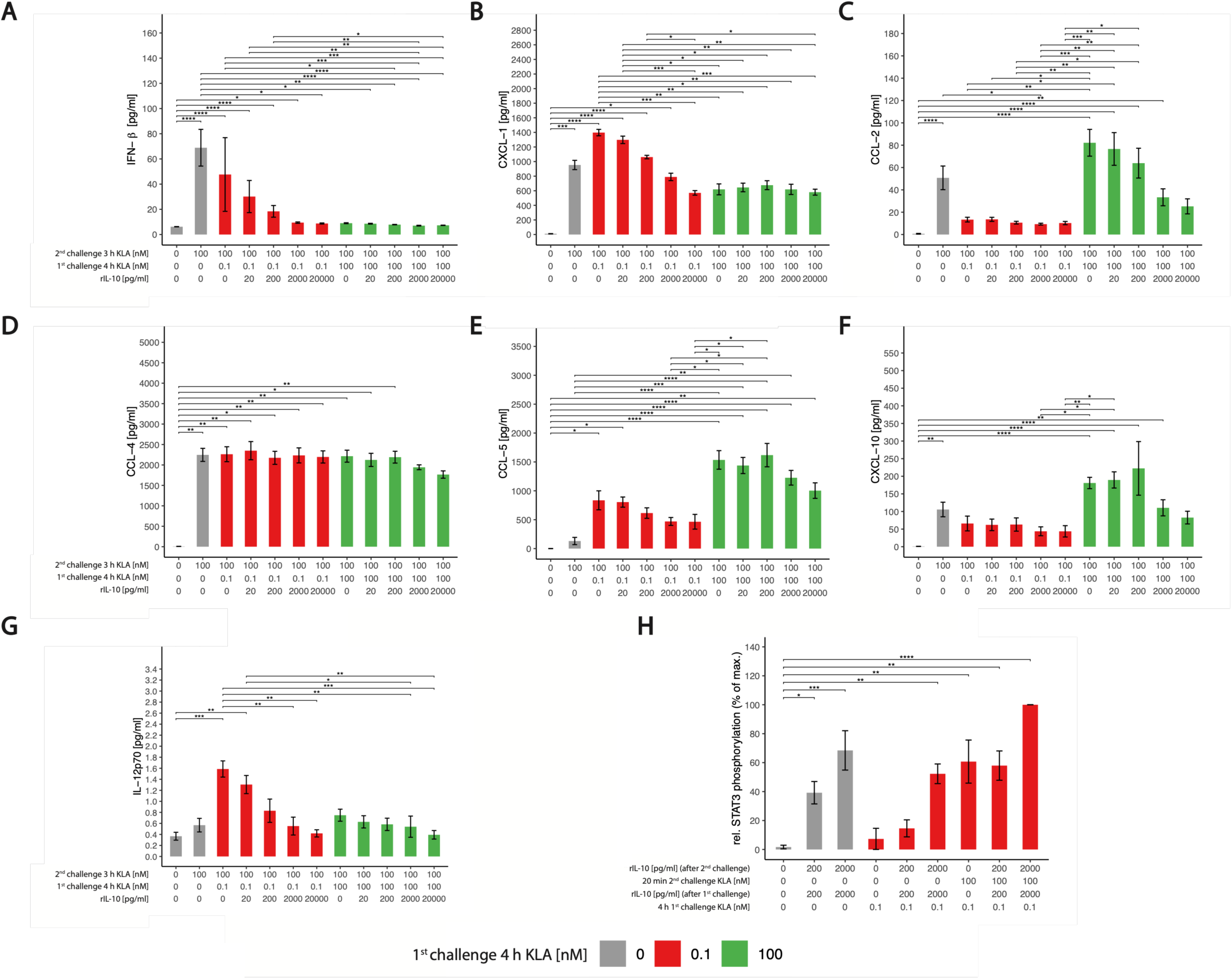
IL-10 without strong prestimulation is insufficient to induce hyporesponsiveness. BMDMs were incubated with 5% FBS DMEM. During primary stimulation, cells were treated with 0, 0.1 or 100 nM KLA. 20 minutes later, recombinant murine IL-10 was added in depicted concentrations. After 4 hrs, cells were washed, medium was replaced with 5% FBS DMEM and cells were incubated for 1 hr. Subsequently, restimulation was performed using 0 or 100 nM KLA. 20 minutes later, recombinant murine IL-10 was added in depicted concentrations. After 3 hrs, supernatants were collected, processed with the LegendPlex™ Multiplex Assay Kits and cytokine and chemokine levels for (A) IFN-b, (B) CXCL-1, (C) CCL-2, (D) CCL-4, (E) CCL-5, (F) CXCL-10, and (G) IL-12p70 were determined by using a LSRII flow cytometer (BDBiosciences). Raw data was processed with LegendPlex™ Desktop software. Each replicate was performed with BMDMs from different mice and data sets include 6 replicates (n=6). Data are given as mean ± standard error of mean. Kruskal–Wallis test with post-hoc Dunn-Bonferroni comparisons: \**p* < .05, \*\**p* < .01, \*\*\**p* < .001, \*\*\*\**p* < .0001. **(H) Recombinant IL-10 induces STAT3 phosphorylation.** BMDMs were incubated with 5% FBS DMEM. During primary stimulation, cells were treated with 0, 0.1 or 100 nM KLA. 20 minutes later, recombinant murine IL-10 was added in depicted concentrations. After 4 hrs, cells were washed, medium was replaced with 5% FBS DMEM and cells were incubated for 1 hr. Subsequently, restimulation was performed using 0 or 100 nM KLA. 20 minutes later, recombinant murine IL-10 was added in depicted concentrations for 20 minutes. Subsequently, cells were washed, fixed, permeabilized and intracellularly stained for phosphorylated STAT3. Median fluorescence intensity of samples was determined by using a LSRII flow cytometer (BDBiosciences). Raw data was processed with FlowJo™. Data is given in % of maximal median fluorescence intensity within each replicate (set as 100%) and was normalized by subtracting median fluorescence intensity of the sample with the detected minimal median fluorescence intensity (set as 0%). Each replicate was performed with BMDMs from different mice and data set includes 3 replicates (n=3). Data are given as mean ± standard error of mean. Kruskal–Wallis test with post-hoc Dunn-Bonferroni comparisons of each sample with unstimulated control (planned comparisons): \**p* < .05, \*\**p* < .01, \*\*\**p* < .001, \*\*\*\**p* < .0001.

**Fig. S6:**
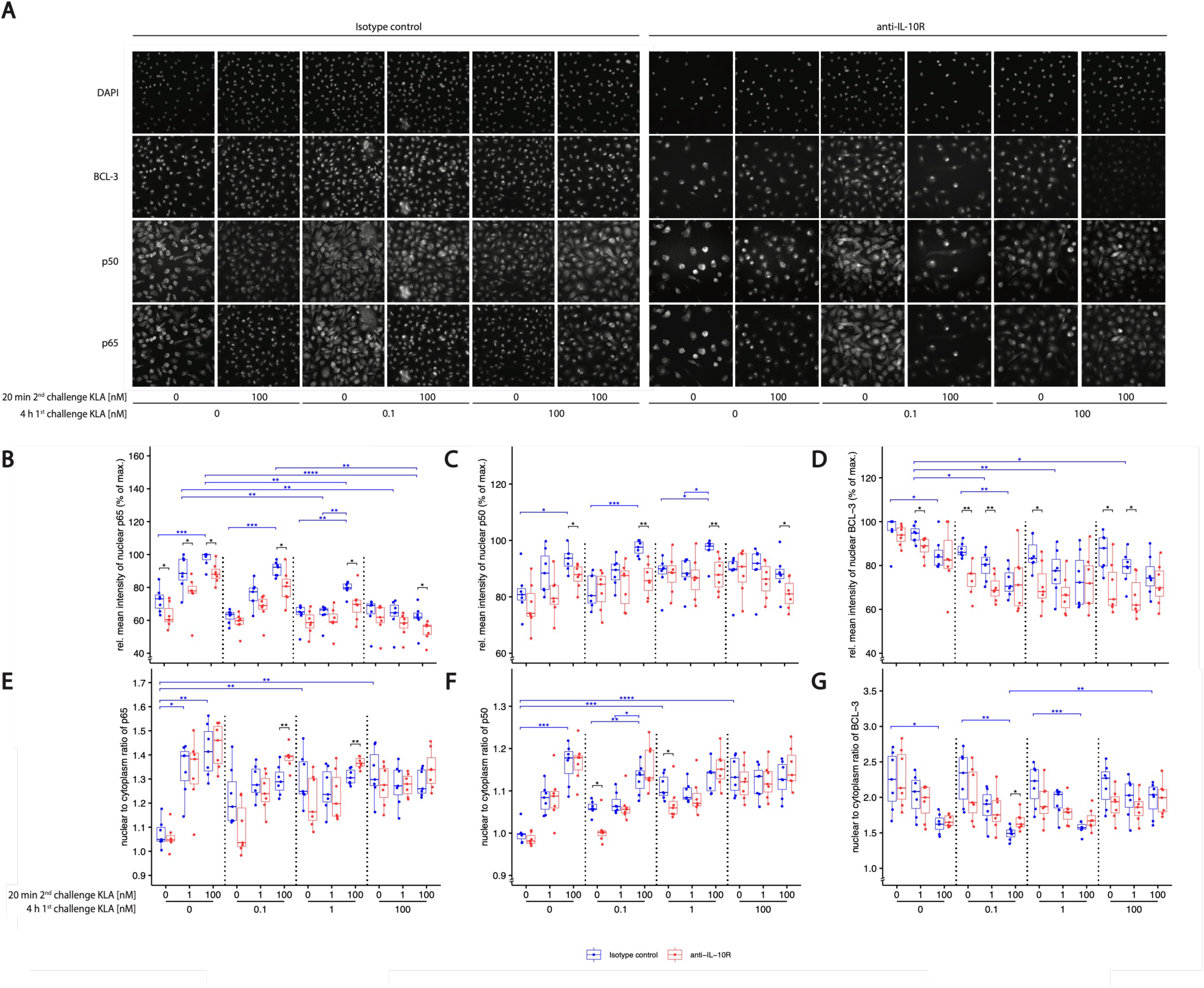
Fluorescence microscopy images of p65, p50 and BCL-3 localization and ratio nuclear vs. cytoplasmic localization. (A) BMDMs were incubated with 5% FBS DMEM containing either IL-10R blocking antibody or its isotype control. During primary stimulation, cells were treated with 0, 0.1, 1 or 100 nM KLA. After 4 hrs, cells were washed, and medium was replaced with 5% FBS DMEM containing either IL-10R blocking antibody or its isotype control and cells were incubated for 1 hr. Subsequently, restimulation was performed using 0, 1 or 100 nM KLA. After 20 mins, cells were washed, fixed, permeabilized and intracellularly stained for p65, p50, and BCL-3, as well as stained with DAPI, and images were acquired using the CellInsight CX7 Pro HCS Platform (Thermo Fisher Scientific) equipped with an 40x objective lens. (B-G) Raw data was processed with CellProfiler™. Data is given in relative mean nuclear intensity as % of maximal nuclear fluorescence intensity within each replicate (set as 100%) (B-D) or nuclear to cytoplasmic ratio (E-G) of the corresponding fluorophore-coupled antibody for p65, p50 or BCL-3. Each replicate was performed with BMDMs from different mice and data sets include at least 6 replicates (n=6). Kruskal–Wallis test with post-hoc Dunn-Bonferroni comparisons: \**p* < .05, \*\**p* < .01, \*\*\**p* < .001, \*\*\*\**p* < .0001.

**Fig. S7:**
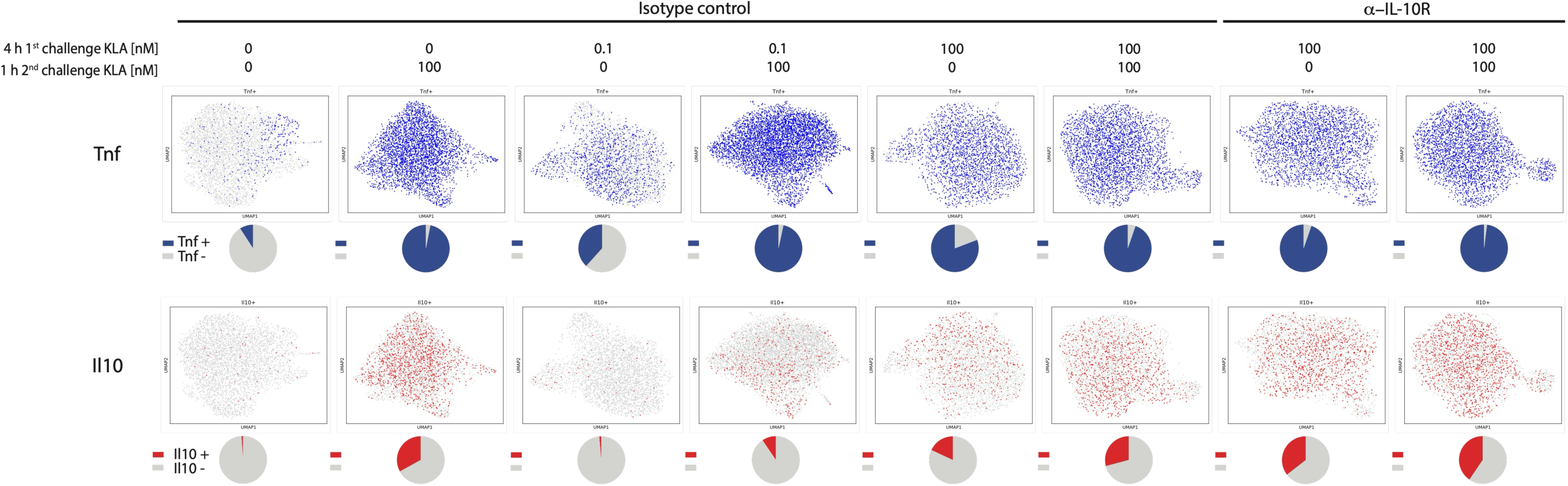
Single cell sequencing of Tnf and Il10 gene expression. BMDM were incubated with 5% FBS DMEM containing either IL-10R blocking antibody or its isotype control. During priming, cells were stimulated with 0, 0.1, or 100 nM KLA. After 4 hrs, cells were washed, and medium was replaced with 5% FBS DMEM containing either IL-10R blocking antibody or its isotype control and cells were incubated for 1 hr. Subsequently, re-stimulation was performed using 0 or 100 nM KLA. After 1 hr, cells were collected and subjected to single cell RNA sequencing. Pre-processed data was used for unsupervised UMAP clustering of all treatment conditions. Each dot represents one cell either negative (grey) or positive (color) for the respective expressed gene. Percent of expressing and non-expressing cells depicted as pie charts.

**Fig. S8:**
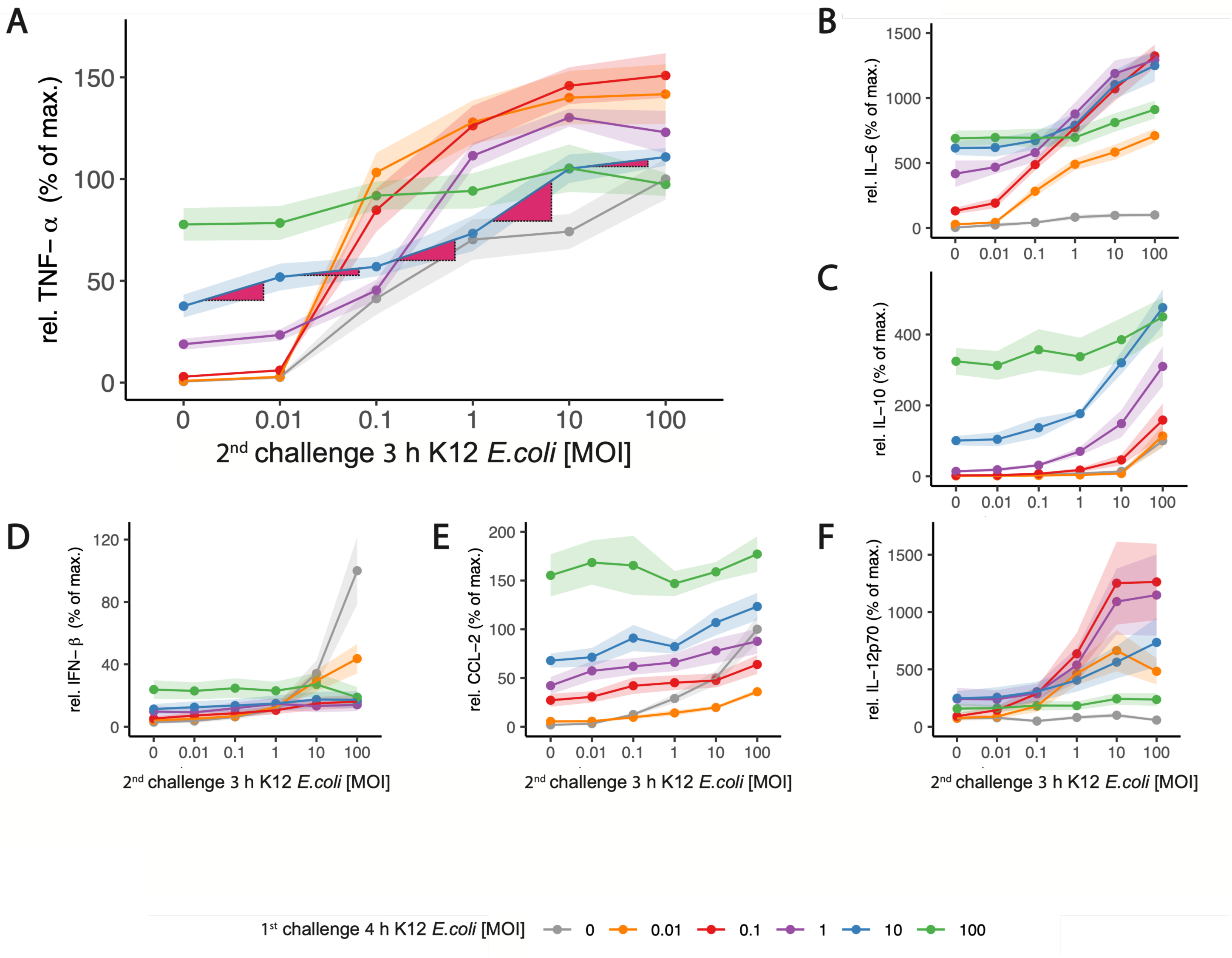
Bacterial load during 1^st^ challenge sets TNF-α and IL-6 threshold bacterial load for increased responsiveness toward 2^nd^ challenge. During primary and secondary stimulation, BMDMs were incubated with 5% FBS DMEM. During the 1^st^challenge, cells were stimulated with 0, 0.01, 0.1,1, 10, and 100 MOI of heat-inactivated K12 *E.coli*. Restimulation was performed using 0, 0.01, 0.1,1, 10, and 100 nM MOI of heat-inactivated K12 *E.coli*. After 3 hrs, supernatants were collected, processed with the LegendPlex™ Multiplex Assay Kits and cytokine and chemokine levels of (A) TNF-α, (B) IL-6, (C) IL-10, (D) IFN-β, (E) CCL-2, and (F) IL-12p70 were determined. Raw data was processed with LegendPlex™ Desktop software. Each replicate was performed with BMDMs from different mice. Data include at least 6 replicates (n=6). Data was normalized to the mean maximal cytokine secretion induced by restimulation in naïve (0 MOI primary challenge) cells (set as 100%). Data are given as mean ± SEM. P-values: Kruskal–Wallis test with post-hoc Dunn-Bonferroni comparisons. In (A), Dark red-colored triangles attached to blue curve (1^st^ challenge = 10 MOI) indicate the increase in TNF-α secretion following 2^nd^ stimulation for every 10-fold increase in bacterial load of the secondary challenge. The biggest increase can be seen between the 2^nd^ challenges with 1 and 10 MOI. Similarly, for a 1^st^ challenge 0.1 or 1 MOI, the strongest changes in the responses can be seen for 2^nd^ challenges between 0.01 and 0.1 or 0.1 and 1 MOI, respectively. The same behavior (strongest increases following secondary stimulation matching the first) was observed for IL-6. Stimulation with a MOI of 100 resulted in persistent production of TNF-α and IL-6 and hypo-responsiveness toward secondary challenge.

## Supplementary Text 1: A minimal model of IL-10 licensing and TLR4 memory during a macrophage response to infection

### 1. Model purpose and scope

This model illustrates the functional consequence of two coupled features of macrophage regulation, namely history-dependent IL-10 licensing and quantitative (match-or-exceed) memory, over the course of a single infection. It is deliberately minimal and dimensionless: it is not fitted to kinetic data, but is used to compare three regulatory logics under identical parameters, so that any difference in outcome is attributable to the single per-cell term by which IL-10 acts on inflammatory output.

### 2. State variables

To encode the stimulation-specific memory of macrophages, the model uses a series of K 1 hierarchical bins (rungs), representing geometrically spaced levels of TLR4 experience. So the base model (with K = 12) integrates 13 macrophage states plus four scalar entities. The resolution extension adds a transition of to an M2-like state (Section 10):

- P(t)— pathogen load, normalized to its carrying capacity K_p_;
- M_j_(t), j = 0, …, K — macrophages occupying memory rung j (a discretized adaptation set-point, see below), with M_0_ the naive cells;
- E(t) — lumped pro-inflammatory output (a TNF-*α*/IL-6 proxy) that drives pathogen clearance;
- D t — tissue-damage proxy;
- M2(t) — pro-resolving (M2) macrophages (added in the extension, Section 10).

### 3. Sensing and history variables

**Sensing drive.** All cells sense the same drive d t, a saturating sum of a pathogen term and a sterile damage-associated (DAMP) term, so that released damage re-activates the same TLR4-type pathway. The drive is a dynamic quantity computed from the current state (it is not a free parameter and does not appear in the parameter Table 1); it enters the escape gate, the inflammatory output, and IL-10 production below:

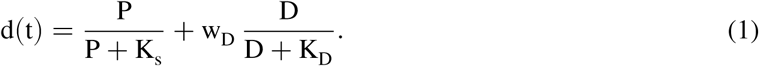

**IL-10 suppression factor.** The fraction of pro-inflammatory output that remains under the current IL-10 level (a Hill function), with f = 1 when IL-10 is absent or its receptor is blocked and f → 0 at saturating IL-10:

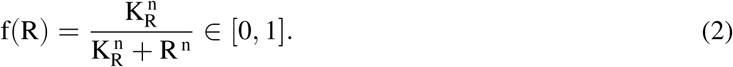

**Memory set-points (the ladder).** Cells carry a discretized adaptation set-point on a fixed geometric ladder spanning the sensing dynamic range; the ladder positions are set by that range, not fitted:

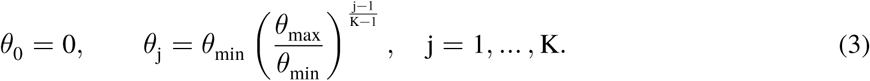

**Licensing.** A cell’s susceptibility to IL-10 is a smooth (graded) function of its memory set-point: only cells with a strong stimulation history are IL-10-suppressible:

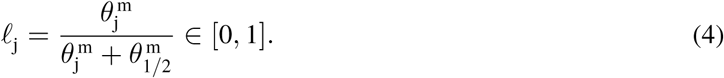

**Escape (match-or-exceed).** A stimulus that exceeds a cell’s set-point re-activates it. This soft threshold also drives the upward ratchet of the ladder:

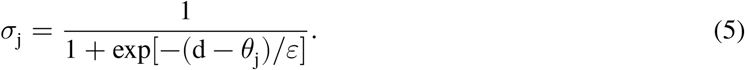

## 4. Dynamics

**Macrophage recruitment and memory ratchet.** Cells are recruited at the lowest rung with a baseline term plus an inflammation-driven term, bounded by a finite pool M_max_ and ratchet upward through rungs they exceed, at rate *α* (see Fig. S10).

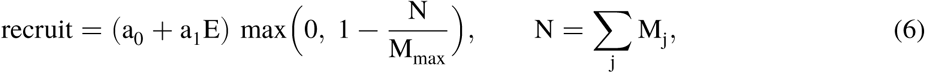

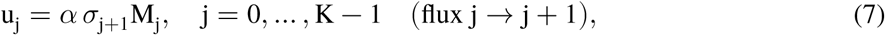

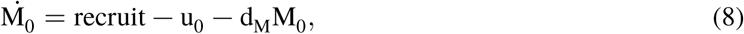

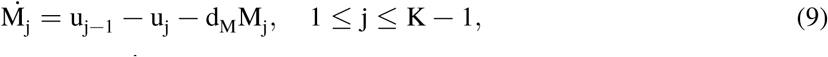

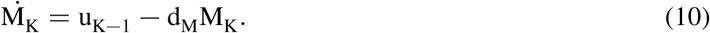

A cohort therefore accumulates at the rung matching the highest drive it has experienced (its memory), and falls silent on a plateau or a falling input.

**Pro-inflammatory output (the only term that differs between the three variants).** The graded per-cell IL-10 amplitude factor is _j =_ 1 − *l*_j_(1 −f) . So μ_j_ = 1 for an unlicensed cell and _j_ f for a fully licensed cell. The population produces inflammatory output in proportion to the common sensing drive d (Eq. 1), weighted by a per-cell output bracket B_j_:

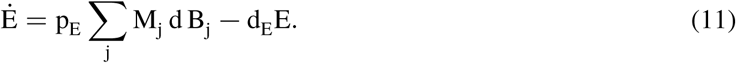

The three variants differ only in B_j_:

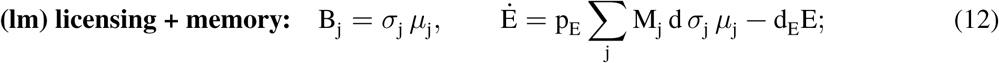

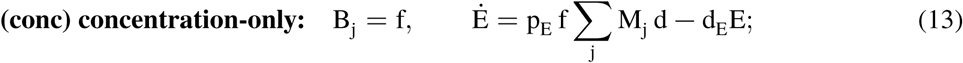

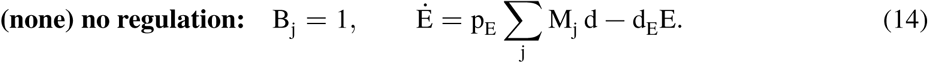

Under history-gated licensing, a cell contributes only when its escape gate is open (σ_j_), and IL-10 then scales the amplitude of that firing by _j_, to the extent the cell is licensed. This means: memory sets *which and how many* cells fire, IL-10 sets *how strongly* each firing cell outputs, and the two act multiplicatively (Section 6).

**IL-10, variable R** (identical in all three variants). Production is driven by the same sensing drive d (Eq. 1) and weighted by the licensing function. As reported in Fig. S2 (compare prestimulation with 0.3 nM KLA in **A** to **E**), cells begin secreting IL-10 exactly where they become able to respond to it. In addition, there is a small tonic term and a constant ambient influx:

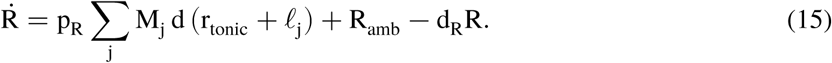

**Pathogen, variable P.** Logistic growth, cleared by a saturable, inflammation-dependent term:

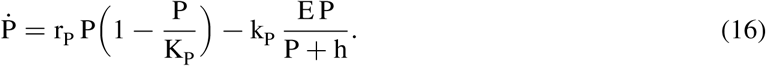

**Tissue damage, variable D.** Driven by inflammation and by pathogen burden, repaired at a constant rate:

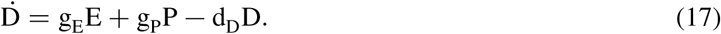

### 5. The three regulatory variants

All three variants we are exploring use identical parameters for pathogen kinetics, macrophage recruitment, and IL-10 production. The variants differ only in which cells IL-10 suppresses.

- **Licensing + memory (lm).** Only history-licensed cells (*lj* >0) are IL-10-suppressible, and the escape gate _j_ lets any cell respond to a stimulus exceeding its remembered level. Newly recruited (naive) cells are refractory to ambient IL-10.
- **Concentration-only (conc).** Every cell’s output is scaled by f R regardless of history: ambient IL-10 pre-mutes cells in proportion to its concentration.
- **No regulation (none).** IL-10 has no effect on output (f ≡ 1); pure pathogen/DAMP sensing.

### 6. How single-cell data determine the shape of the modeled inflammatory responses

How memory (σ_j_) and IL-10/licensing (σ_j_, f) combine in the per-cell bracket B_j_ is a modeling choice that strongly affects the dynamics. We considered three functional forms and discriminated among them using single-cell TNF measurements (see Fig. 2 H, I in the main text).

**Candidate forms.** One potential way to implement the influence of TLR4 memory and IL-10 is an *escape-dominant additive* bracket, in which an escaping (firing) cell contributes its full output and only the non-escaping fraction is IL-10-suppressible (so a firing cell is *immune* to IL-10), with an IL-10-independent residual *γ*:

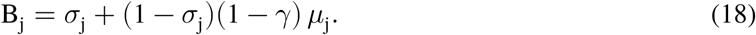

A somewhat simpler *threshold-shift* alternative folds IL-10 into the gate itself: IL-10 raises a licensed cell’s effective set-point rather than reducing its amplitude. This means that one sigmoid carries both controls, and because *σ*_j_ also drives the ratchet, IL-10 would couple to memory *formation*:

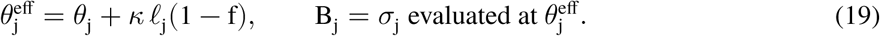

The adopted *product* form (Eq. 12) instead has IL-10 multiply the output of firing cells irrespective of escape.

The three forms differ on one experimentally accessible question: *is a cell that is already firing (escaped) suppressible by IL-10?*

| form | output bracket $B_j$ | firing cell IL-10-suppressible? | verdict from data |
| --- | --- | --- | --- |
| additive | $\sigma_j + (1 - \sigma_j)(1 - \gamma)\mu_j$ | **no** — escape overrides suppression | contradicted |
| threshold-shift | $\sigma_j$ at $\theta_j^{\text{eff}}$ | indirectly — alters *whether* it fires, not amplitude | not the observed mode |
| **product (adopted)** | $\sigma_j \mu_j$ | **yes** — amplitude scaled by $\mu_j$ | **supported** |

**Single-cell data.** (For details, see description of data in Fig. 2 I, H.) Macrophages were primed (first challenge 0, 1, or 100 nM KLA), rested, and re-challenged (second challenge 0 or 100 nM), with isotype control or anti-IL-10R antibody on the 100 nM-primed arm; per-cell *Tnf* expression was quantified in TNF^+^ cells (n 6 7–9 1 10^3^ per condition). Two axes emerge: *Memory sets the responding fraction*: with increasing first-challenge dose the TNF^+^ fraction rises steeply ( 8 9 38 5 81 4 at second challenge 0), a fraction-encoded response. *IL-10 sets the per-cell amplitude of firing cells*: at the saturating second challenge ( 95 TNF^+^, so the fraction is fixed), the per-cell TNF among firing cells is lower for heavily-primed cells (100/100, mean 2 56) than for lightly-primed cells (1/100, mean 3 72), and IL-10R blockade restores the heavily-primed cells to the lightly-primed level (100/100 + anti-IL-10R, mean 3 59). These are escaped, firing cells, yet IL-10 still suppresses their amplitude. This directly contradicts the additive form’s escape-immunity and identifying amplitude (gain) control (Fig. S9A).

**Uniform gain, not a suppressed subpopulation.** Three tests on the single-cell distributions distinguish a near-uniform per-cell gain reduction from an IL-10-silenced subpopulation. (i) The suppressed distribution (100/100, isotype) is statistically unimodal (according to Hartigan dip test p 0 95) no separated low-TNF mode. (ii) IL-10R blockade lifts the entire distribution by ≈ 1 log unit at every quantile (from 1 16 at the lowest decile to 0 88 at the highest; the quantile–quantile map is close to a constant offset, Fig. S9B,C), so essentially all firing cells are suppressed, not a fraction of them. (iii) Suppression barely changes the spread (standard deviation 0.70 versus 0.64 log units; a two-population mixture would broaden it), and a two-component Gaussian improves barely on a single component here.

Together these observations exclude a discrete IL-10-refractory subpopulation and show that the memory-controlled firing fraction and the IL-10-controlled per-cell amplitude are separable quantities that combine **multiplicatively**. The model therefore represents IL-10 as an amplitude factor 1 1 f acting on firing cells, giving the product output term B_j_ = *σ*_j_ *μ*_j_.

### 7. Initial conditions, integration, and tissue damage metrics

The system starts from rest: P(0)= P_0_; M_0_(0) = M_n0_ with all other rungs zero; E(0) = 0; R 0 = R_amb_/d_R_ (steady-state ambient tone); D(0) = 0; and M2(0) = 0. Equations are integrated to t = 240 h with LSODA (SciPy), relative tolerance 10^−7^, absolute tolerance 10^−10^. The summary metrics are the peak and final P, the final D, and the cumulative tissue insult

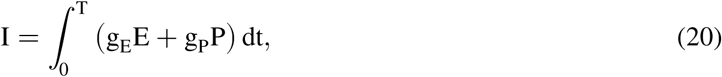

decomposed into an inflammatory burden 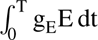 (penalizing failure to resolve) and a pathogen burden 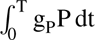 (penalizing failure to clear).

### 8. Parameters

**Table 1.**
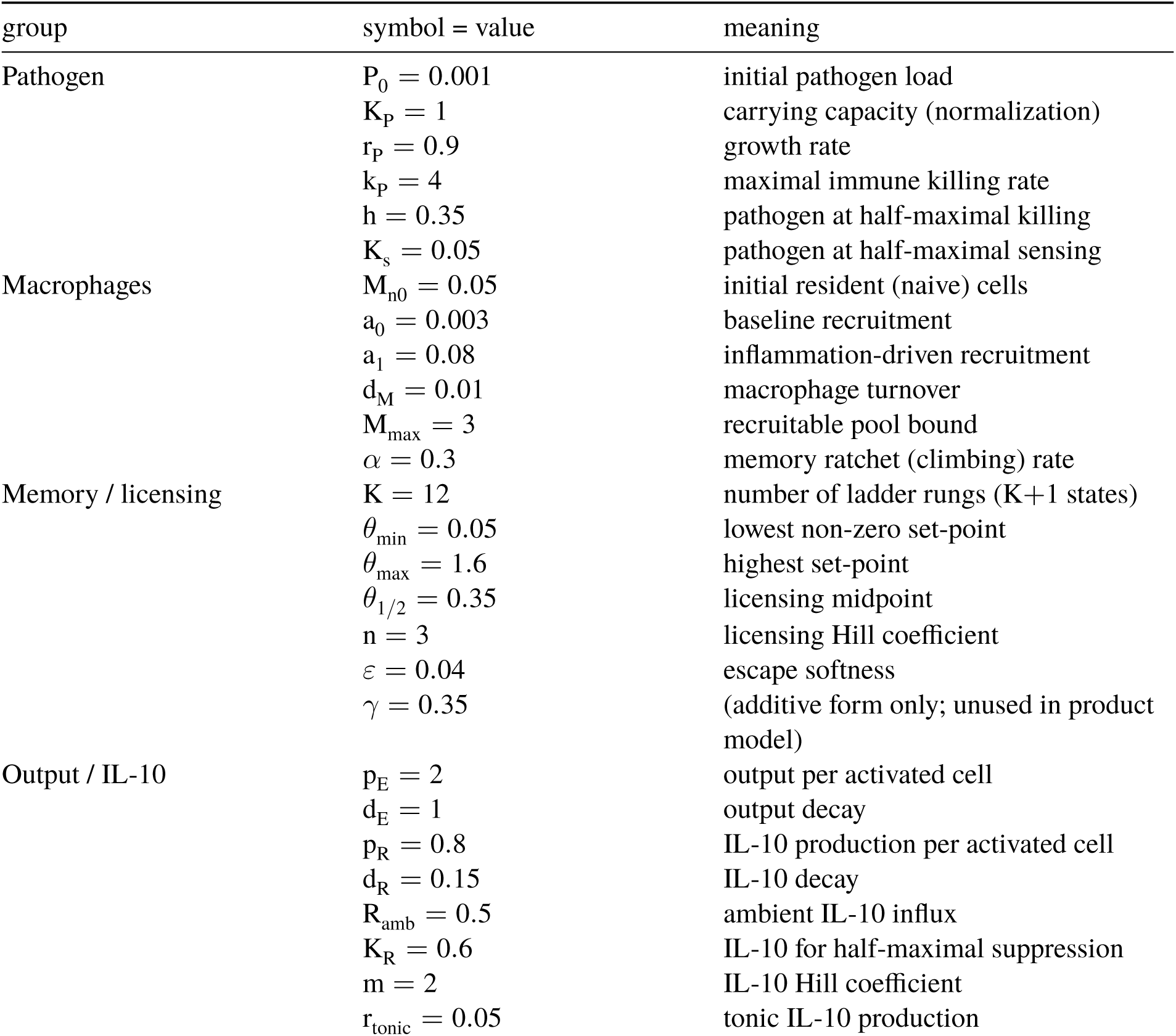
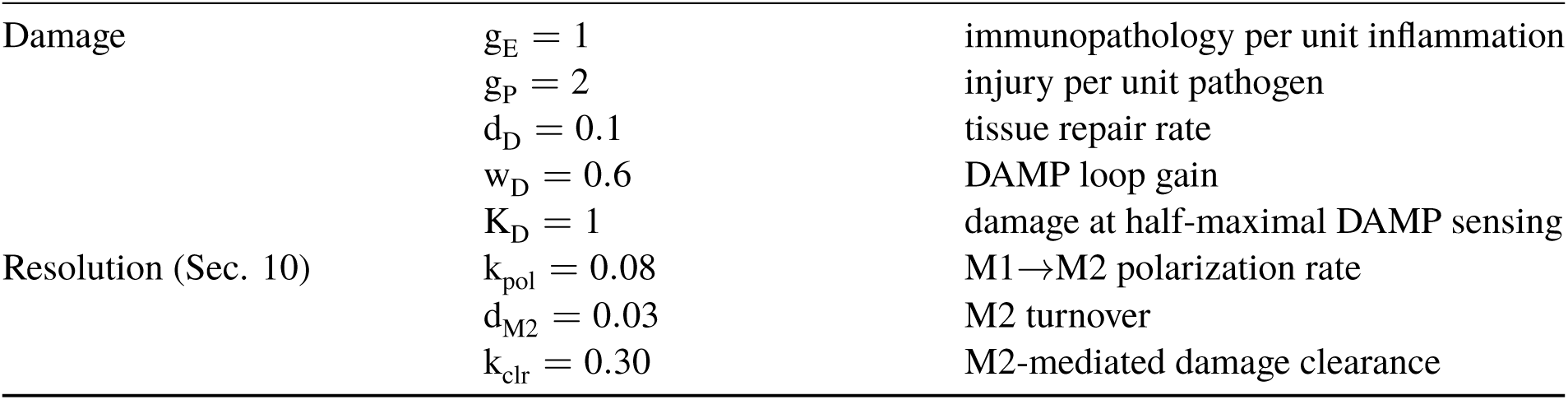
Default parameters are either dimensionless or rates per hour.

### 9. Results at the default operating point

**Table 2.**
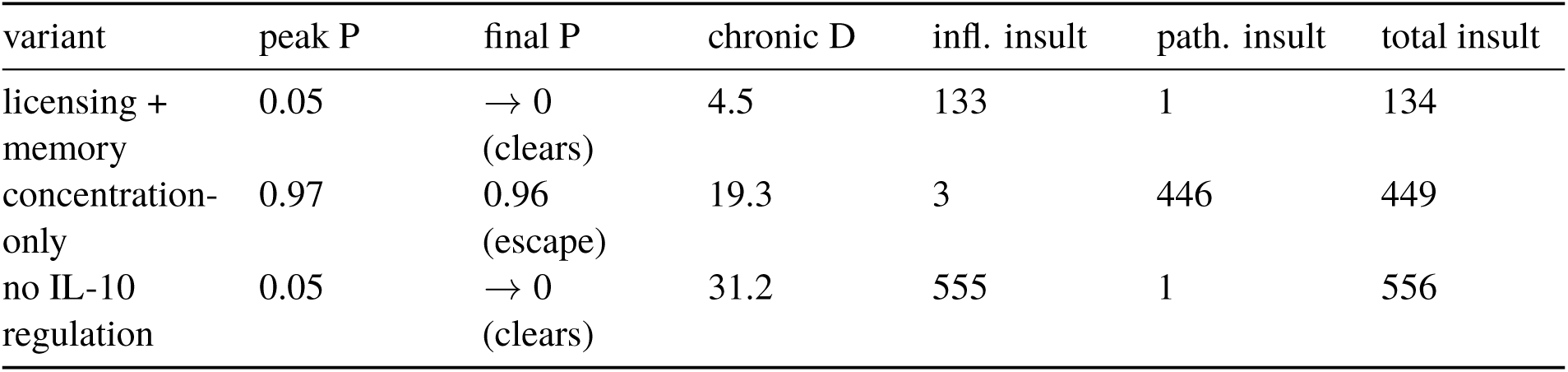
Outcome at the default operating point (base model, product form, t = 240 h).

**Interpretation.** Licensing+memory is the only variant that is low on *both* failure axes (main-text Fig. 5A). Concentration-only pre-mutes naive cells through the ambient IL-10 tone, so the rising pathogen is never met and it escapes (peak P ≈1); its injury is pathogen-driven. No-regulation clears the pathogen but, lacking an IL-10 brake, ignites the DAMP loop into a non-resolving sterile-inflammation state (chronic D ≈ 31). Licensing keeps newly recruited cells refractory to ambient IL-10 so they clear the pathogen early. Then, once licensed, they become IL-10-suppressible, letting IL-10 terminate the loop. A residual low-grade sterile inflammation persists (final D 4 5) because the DAMP plateau is sub-maximal, so cells license only partially and IL-10 suppression is incomplete. This indefinite persistence is a limitation of the base model, which has no active resolution program. The M2 extension below supplies a mechanism for resolution.

### 10. Extension: active resolution by M1→M2 polarization

**Motivation.** The base model’s only brake is IL-10, and it is partial; there is no efferocytosis, no M1→M2 switch, no pro-resolving-mediator program. The bistable DAMP loop therefore has no exit and sterile inflammation persists at a fixed point indefinitely, which is unrealistic for an infection cleared within a day. Therefore, we add a minimal, mechanistic resolution arm in which the most adapted, most IL-10-suppressed macrophages convert to an M2/pro-resolving phenotype, so that resolution becomes part of the same IL-10/licensing program.

**Polarization flux.** A new pool M2 receives cells from the ladder. The per-rung polarization weight is the IL-10 suppression that the adapted cell at rung j experiences and is the product of its licensing and the active IL-10 fraction:

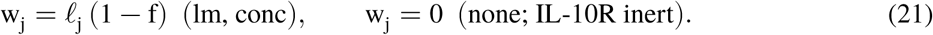

Highly-licensed cells (*l*_j→_1) under strong IL-10 (f→0) convert fastest. The conservative flux q_j =_ k_pol_ w_j_ M_j_ is removed from each ladder equation and summed into the M2 pool, which clears the DAMP source (note additional term in equation (23)).

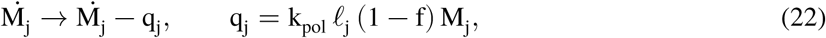

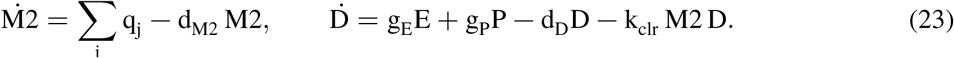

M2 cells produce no inflammatory output. Having left the ladder, they are absent from the ∑_j_ M_j_ that drives ^Ė^. The rates k_pol_, d_M2_, k_clr_ are illustrative, not fitted; k_pol_ = 0 recovers the base model.

**Table 3.** Outcome with active resolution (product form, t = 240 h; time courses in main-text Fig. 5B).

| variant | peak P | final P | chronic D | max M2 | total insult |
| --- | --- | --- | --- | --- | --- |
| licensing +<br>memory | 0.05 | $\rightarrow 0$ (clears) | 0.0 (resolves) | 0.52 | 60 |
| concentration-only | 0.97 | 0.97 (escape) | 15.7 | 0.08 | 453 |
| no IL-10<br>regulation | 0.05 | $\rightarrow 0$ (clears) | 31.2 | 0.00 | 556 |

**Note on a non-physiological simplification in the model:** Setting $w_j = 0$ under no-regulation makes resolution purely IL-10-driven, so that variant cannot resolve at all. Real M2 polarization also has IL-10-independent drivers (IL-4/IL-13, efferocytosis, TGF- $\beta$ ); a small IL-10-independent term would let no-regulation partially resolve and could reorder it against concentration-only.

**Note on a non-physiological simplification in the model.** Setting w_j_ = 0 under no-regulation makes resolution purely IL-10-driven, so that variant cannot resolve at all. Real M2 polarization also has IL-10-independent drivers (IL-4/IL-13, efferocytosis, TGF-*β*); a small IL-10-independent term would let no-regulation partially resolve and could reorder it against concentration-only.

**Fig. S10** shows how the memory rungs of the model with the resolution extension are filled over time and then lose cells again as these enter the resolution M2-like population. The levels in the legend correspond to the discretized memory setpoints in Eq. 3. With the M2 extension, higher memory rungs are filled only weakly, as cells escape into the M2 state and inflammation is terminated around 200 hours. In the base model, without M2-like state, inflammation would continue beyond 200 hours (compare Fig. 5A to Fig. 5B) and cells would linger in medium-high memory rungs.

### 11. Global sensitivity of the licensing advantage

**Setup.** We quantify the robustness of the licensing+memory advantage through the two cumulative-insult differences

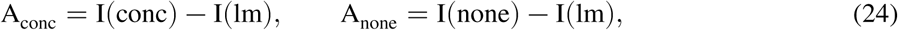

the insult that licensing+memory avoids relative to concentration-only and to no-regulation; the per-draw decomposition into pathogen and inflammatory burden is shown in main-text Fig. 5C. We draw Latin-hypercube samples (N = 300) with every non-structural parameter uniform on 0 5 1 5 its default (structural quantities K_P_, K, the ladder bounds, P_0_, M_n0_, the Hill exponent n, and the unused remain fixed), and rank each parameter’s monotone influence by partial rank correlation (PRCC).

**Base model.** Licensing+memory has the lowest cumulative insult in 80% of draws (beating concentration-only in 83%, no-regulation in 97%; median A_conc_ =306, A_none_ =333). The two advantages behave oppositely, and the PRCC explain why through a sign flip (Fig. S11): the inflammation-cost parameters (output per cell p_E_, DAMP gain w_D_, immunopathology g_E_, the macrophage ceiling M_max_) lower A_conc_ but raise A_none_. Making inflammation expensive penalizes licensing+memory’s own (controlled) response relative to concentration-only’s near-silence, yet simultaneously makes the IL-10 brake more valuable relative to the unbraked case. The pathogen-danger parameter g_P_ acts almost only on A_conc_, punishing concentration-only’s escape. This is the well-known resistance-versus-tolerance trade-off, recovered mechanistically: regulated resistance (licensing+memory) beats indiscriminate suppression when the pathogen is dangerous (g_P_ high) and inflammation is containable (d_E_ high, p_E_/w_D_/g_E_ low); when inflammation is extremely costly and the pathogen benign, tolerating it can win.

**With M1→M2 conversion.** Repeating the analysis on the resolution model increases the advantage of the licensing+memory variant: licensing+memory is now best in 87% of samples (versus 80%), beats concentration-only in 91% (versus 83%), and the no-regulation advantage holds at 96%; the median advantages rise (A_conc_: 306 318, A_none_: 333 387). The mechanism is visible in the PRCC: the inflammation-cost parameters that erode A_conc_ in the base model have weaker influence once M2 is present, because licensing+memory can now resolve the inflammation it generates, so the regime in which tolerance beats resistance contracts. The per-cell output rate p_E_ enters the two advantages with opposite sign, narrowing the clearance gap against concentration-only but widening the resolution gap against no-regulation.

**Caveats.** The analysis uses 50 uniform ranges and PRCC, which assumes monotone parameter effects; a few affinities (e.g. K_R_) may be non-monotone and have their influence understated. The M2 rates are illustrative; their own influence is modest, so the robustness gain is a structural consequence of possessing a resolution arm, not a product of tuning unfitted parameters. Parameters were not refitted to the product-form operating point, so the qualitative ordering and the licensing advantage are robust while absolute magnitudes are not determined by the data or the model variants.

### 12. A parameter set without runaway pathogen in the concentration-only case

Concentration-only regulation fails in most of the parameter space by runaway pathogen escape. However, when the pathogen is intrinsically self-limiting (lower carrying capacity K_P_) and IL-10 suppression is weaker (higher K_R_), the muted response is still sufficient to hold the pathogen at a sub-maximal plateau rather than losing control entirely (Fig. S13; final P ≈0.51, about 73% of ceiling, at K 0 7, K 1 25).

This regime is rather special: across a 50 sweep of the non-structural parameters, concentration-only escapes outright (>85% of ceiling) in ≈ 89% of draws, controls only partially in ≈7, and clears in ≈ 4. Fig. S14 shows where this partial-control regime is located in the (K_R_ K_P_) plane: it is reached by simultaneously weakening suppression and lowering the ceiling from the default operating point (final P 0 96 0 51). The plot also shows that history-gated licensing still incurs the lower cumulative insult there. Partial control therefore softens the concentration-only failure without reversing the ordering: even where blanket suppression avoids runaway pathogen, it trades escape for a persistent sub-maximal pathogen burden and the accompanying damage. As a result, it pays more in total tissue insult than history-gated licensing. The licensing advantage is thus not an artifact of comparing it to an always catastrophically failing scenario with runaway pathogen. It holds whether concentration-only fails by escape or merely by excess injury under partial control.

### Supplementary Figures

**Fig. S9.**
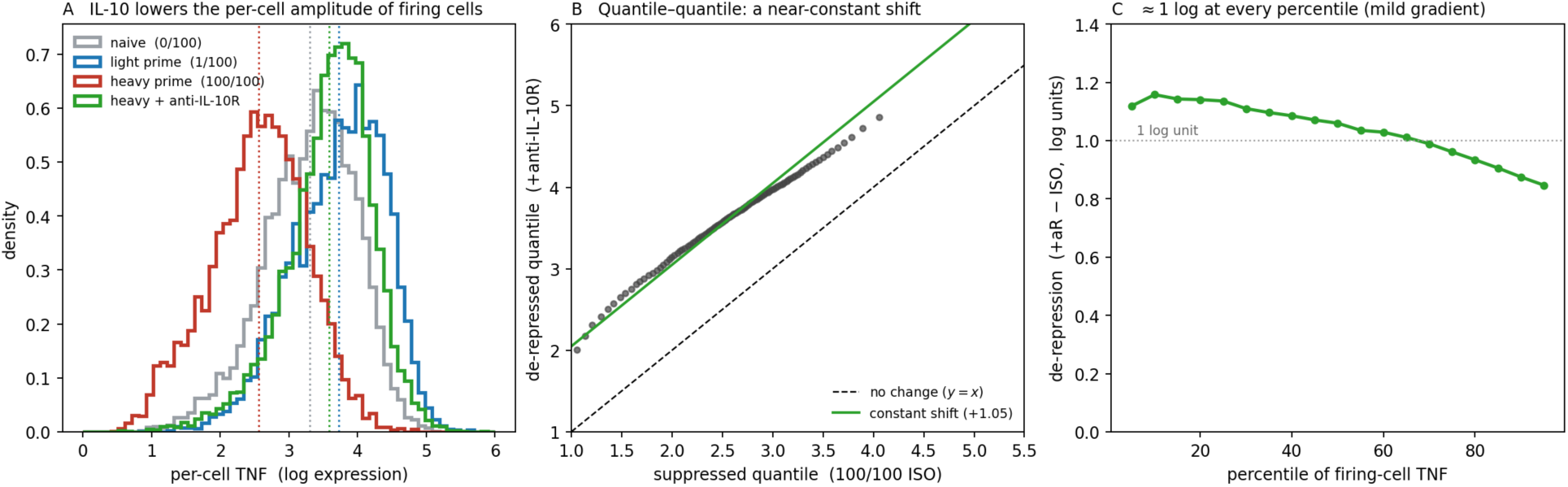
Single-cell TNF in TNF^+^ cells discriminates gain from subpopulation control (data shown in Fig. 2 H, I) **(A)** Per-cell TNF distributions of firing cells: IL-10 slides the heavily-primed distribution (100/100, mean 2.56) downward relative to the lightly-primed one (1/100, mean 3.72), and anti-IL-10R blockade restores it (mean 3.59); naive cells (0/100, mean 3.30) are shown for reference. **(B)** Quantile–quantile comparison of the suppressed (100/100) and de-repressed (+anti-IL-10R) distributions lies close to a constant offset ( 1 05 log units; uniform gain), not peeling away at the lower quantiles as a suppressed subpopulation would. **(C)** The de-repression is ≈ 1 log unit at every percentile, with only a mild gradient (low expressers suppressed slightly more).

**Fig. S10.**
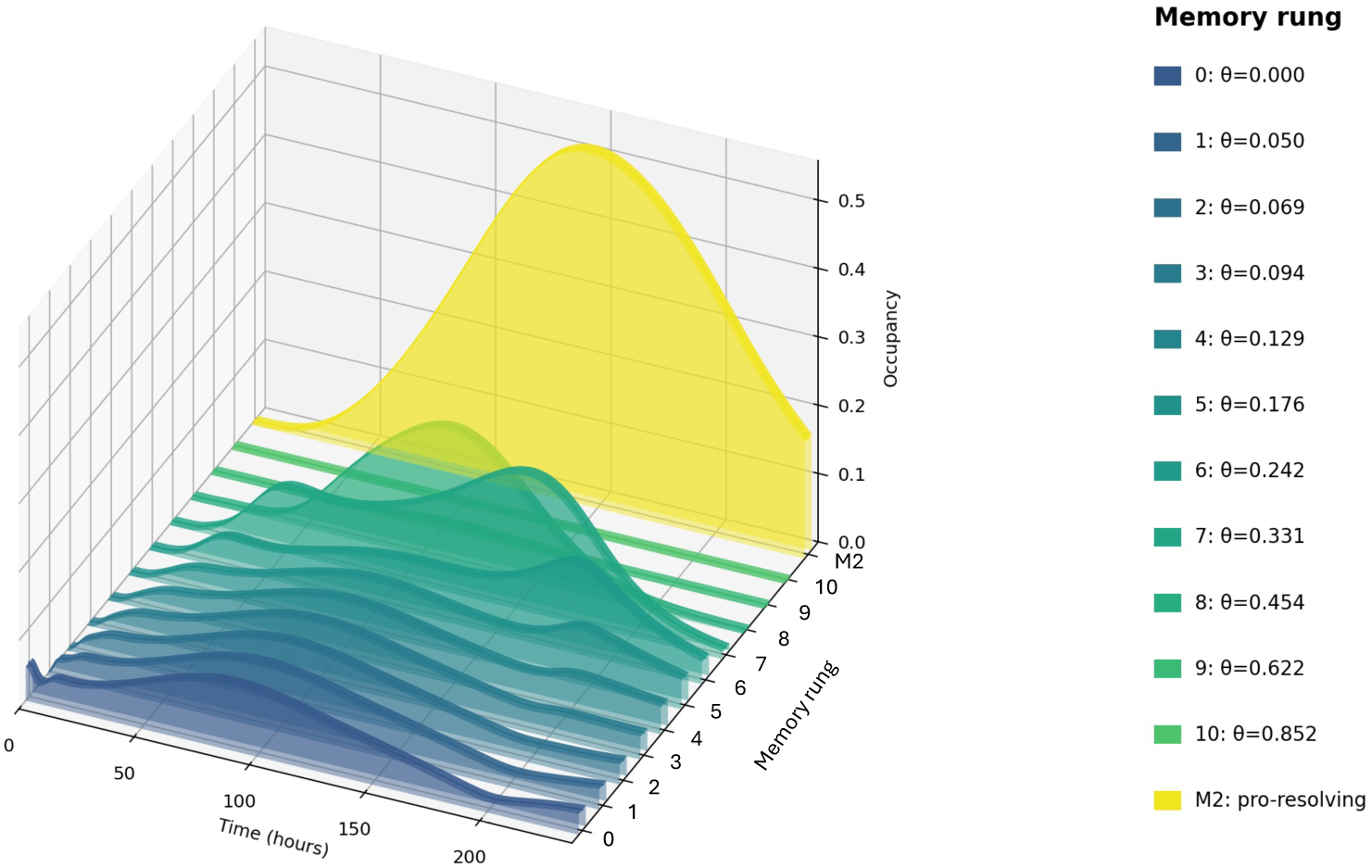
Dynamics of memory rung occupation over time (resolution model). Dynamics of memory rung occupation over time for a simulation of the resolution model. The values in the legend correspond to the memory set points in (Eq. 3). Note that, in this extension with M2-like cells, the higher memory rungs are filled only weakly since the cells escape into the M2 state.

**Fig. S11.**
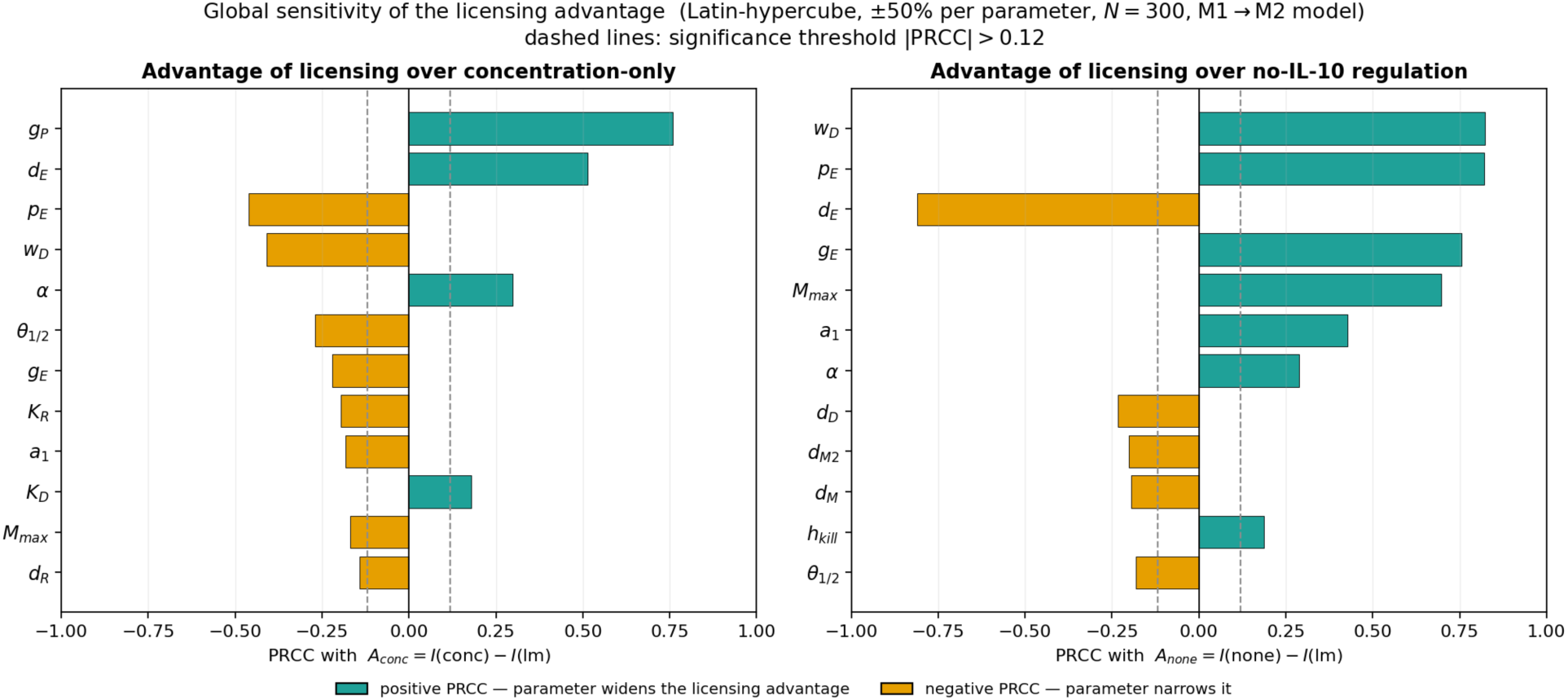
Global sensitivity of the licensing advantage (resolution model). PRCC between each parameter and the advantage of history-gated licensing over **(left)** concentration-only, A_conc_, and **(right)** no regulation, A_none_ (twelve largest-magnitude parameters per panel; teal = widens the advantage, orange = narrows it; dashed lines, PRCC 0 12). The advantage over concentration-only is governed mainly by the pathogen-cost parameter g_P_ and narrowed by weaker IL-10 suppression (larger K_R_); the advantage over no-regulation by inflammation-cost parameters (w_D_, p_E_, g_E_, M_max_). The output rate p_E_ changes sign between the panels, illustrating the resistance/tolerance trade-off in a single parameter.

**Fig. S12.**
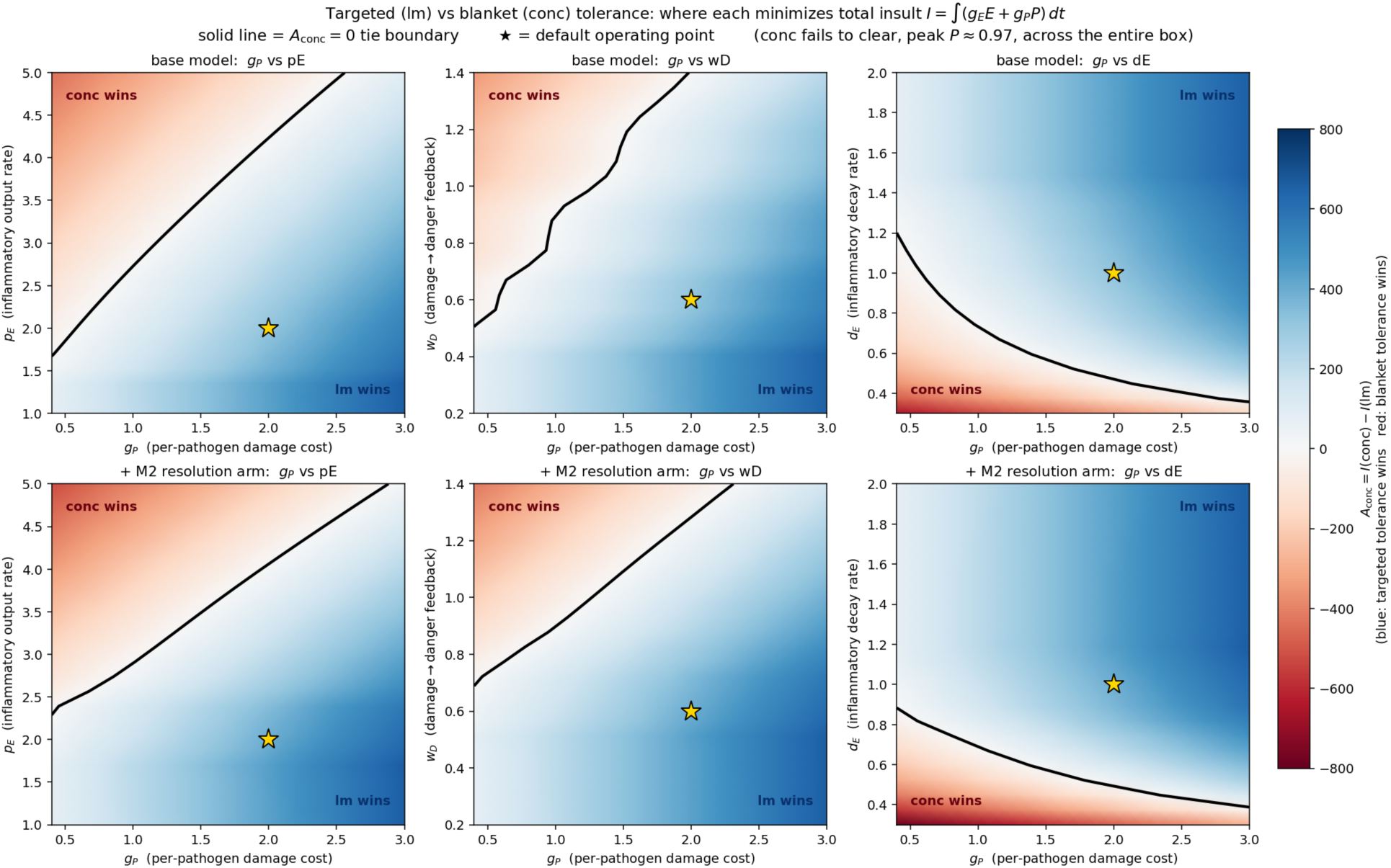
Boundary maps of the licensing advantage. A_conc_ on a 24 × 24 grid (blue: licensing lower insult; red: concentration-only lower; solid line: A_conc_ 0 tie; gold star: default operating point). Columns vary g_P_ against the tolerance parameters p_E_ (left), w_D_ (center), d_E_ (right); top row base model, bottom row with resolution. Licensing gives the lower insult across most of each panel (67–83%), and concentration-only never clears the pathogen anywhere in the box (peak P 0 97): its “wins” are pure tolerance wins. The resolution arm leaves the tie boundary essentially unchanged. Note that the ‘wavy’ structure in the upper middle panel is an artifact, but not of limited sampling, but stemming from the discrete nature of the memory rungs.

**Fig. S13.**
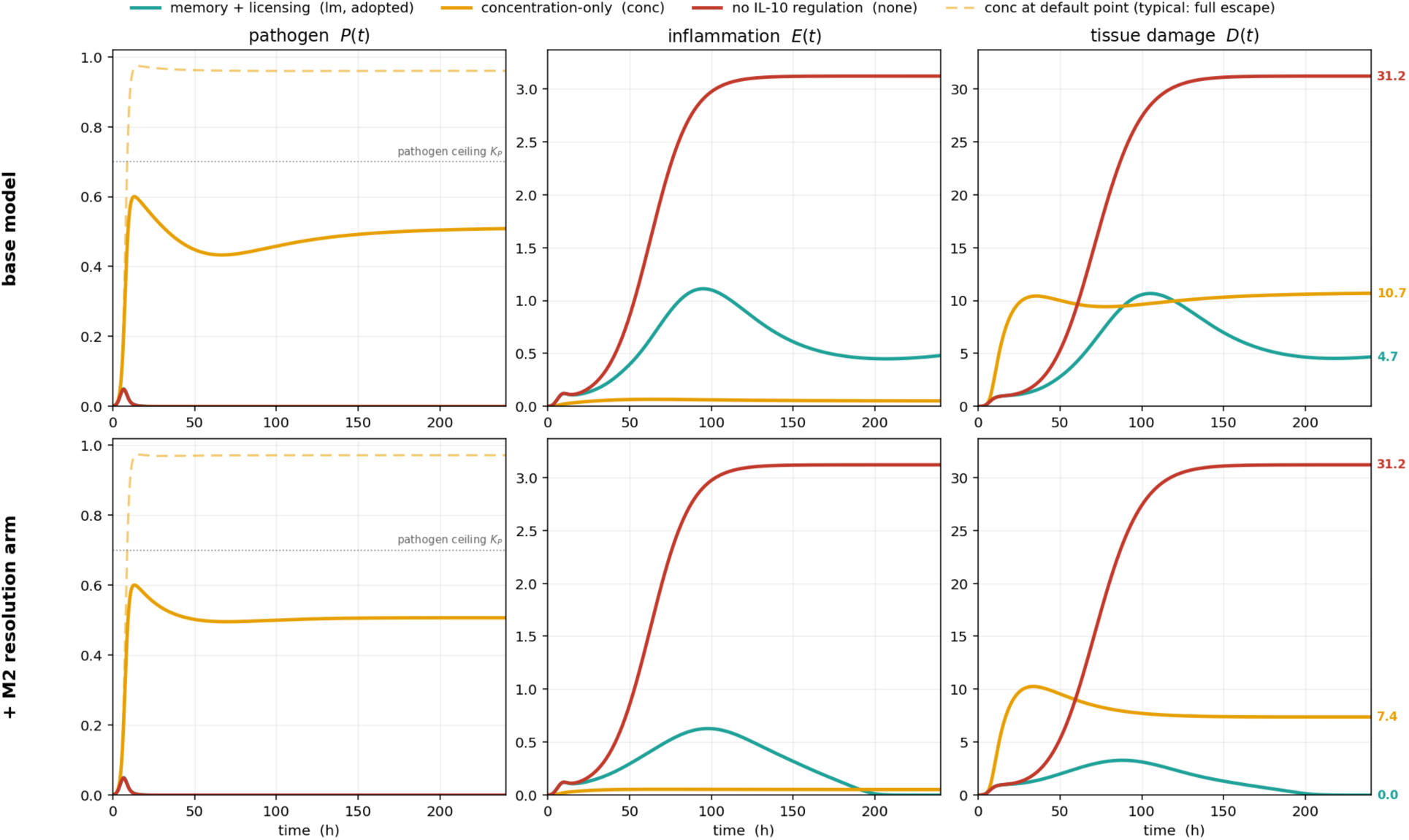
A non-generic operating point where concentration-only fails only partially. Time courses at K_P_ 0 7, K_R_ 1 25 (self-limiting pathogen, modestly weaker IL-10); base model (top) and with resolution (bottom); colors as in Fig. 5A, with the faint dashed curve showing concentration-only at the default point (typical full escape). Here concentration-only holds the pathogen at ≈ 0 51 (≈73 of ceiling) rather than escaping, yet still incurs the greater damage. This regime is non-generic: across a 50 sweep, concentration-only fully escapes (>85 of ceiling) in 89 of draws, partially controls in ≈7%, and clears in ≈ 4%.

**Fig. S14.**
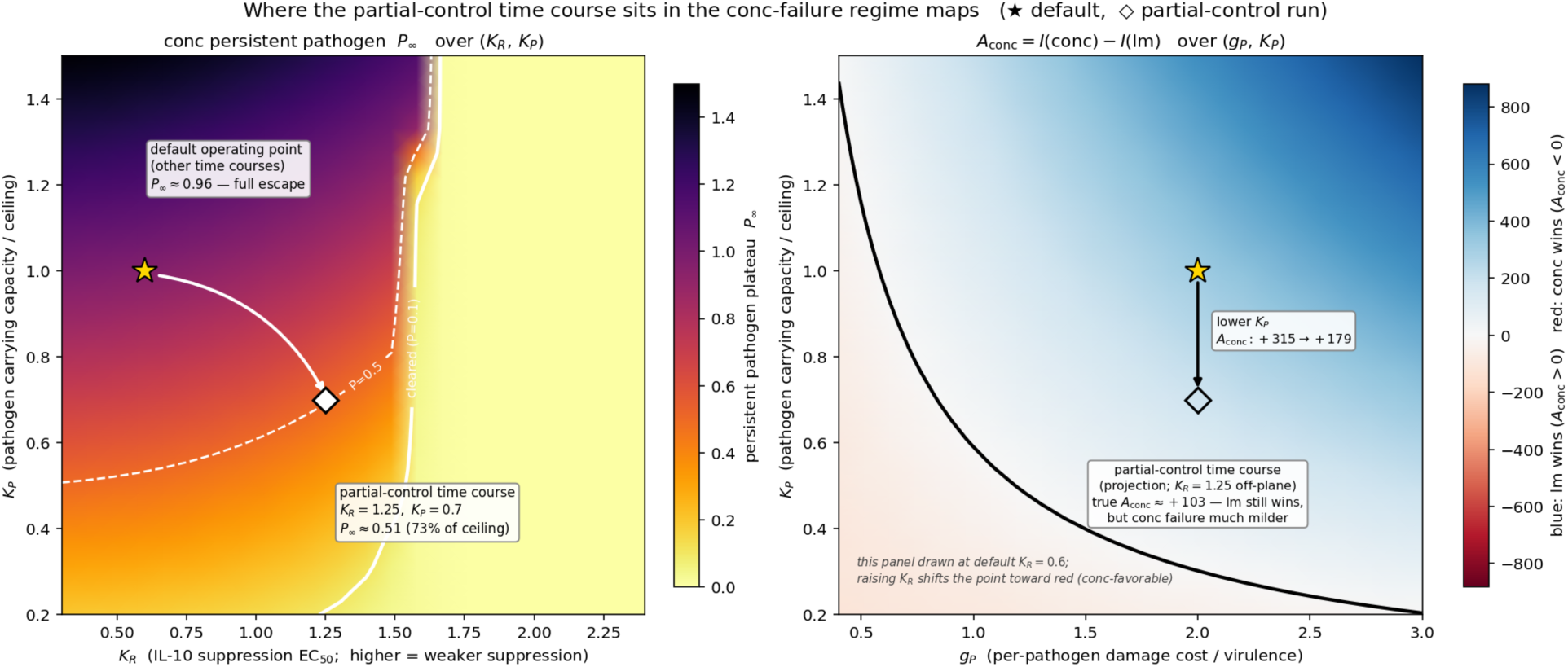
Location of the partial-control regime. (Left) Persistent pathogen plateau P∞ under concentration-only over K_R_, K_P_: the default point (star) escapes fully (P_∞_ ≈0 96); weakening suppression and lowering the ceiling reaches the partial-control point (diamond, P∞≈0 51). **(Right)** A_conc_ over (g_P_ K_P_); because the panel is drawn at the default K_R_, the partial-control point appears as an off-plane projection. Licensing still gives the lower insult there (A_conc_ ≈+103): the partial-control regime makes the concentration-only failure milder without making it the better strategy.

## References

1. K. A. Fitzgerald, J. C. Kagan, Toll-like Receptors and the Control of Immunity. Cell 180, 1044–1066 (2020).

2. S. Akira, K. Takeda, Toll-like receptor signalling. Nat Rev Immunol 4, 499–511 (2004).

3. R. Medzhitov, Toll-like receptors and innate immunity. Nat Rev Immunol 1, 135–145 (2001).

4. T. Kawasaki, T. Kawai, Toll-like receptor signaling pathways. Front Immunol 5, 461 (2014).

5. Y. C. Lu, W. C. Yeh, P. S. Ohashi, LPS/TLR4 signal transduction pathway. Cytokine 42, 145–151 (2008).

6. G. L. Gustafson, M. J. Rhodes, T. Hegel, Monophosphoryl lipid A as a prophylactic for sepsis and septic shock. Prog Clin Biol Res 392, 567–579 (1995).

7. T. Lajqi et al., Training vs. Tolerance: The Yin/Yang of the Innate Immune System. Biomedicines 11, (2023).

8. R. A. Gottschalk et al., Distinct NF-kappaB and MAPK Activation Thresholds Uncouple Steady-State Microbe Sensing from Anti-pathogen Inflammatory Responses. Cell Syst 2, 378–390 (2016).

9. J. J. Muldoon, Y. Chuang, N. Bagheri, J. N. Leonard, Macrophages employ quorum licensing to regulate collective activation. Nat Commun 11, 878 (2020).

10. A. G. Wang et al., Macrophage memory emerges from coordinated transcription factor and chromatin dynamics. Cell Syst 16, 101171 (2025).

11. R. Nakagawa et al., SOCS-1 participates in negative regulation of LPS responses. Immunity 17, 677–687 (2002).

12. F. Nomura et al., Cutting edge: endotoxin tolerance in mouse peritoneal macrophages correlates with down-regulation of surface toll-like receptor 4 expression. J Immunol 164, 3476–3479 (2000).

13. L. M. Sly, M. J. Rauh, J. Kalesnikoff, C. H. Song, G. Krystal, LPS-induced upregulation of SHIP is essential for endotoxin tolerance. Immunity 21, 227–239 (2004).

14. A. Ciesielska, M. Matyjek, K. Kwiatkowska, TLR4 and CD14 trafficking and its influence on LPS-induced pro-inflammatory signaling. Cell Mol Life Sci 78, 1233–1261 (2021).

15. F. Randow et al., Mechanism of endotoxin desensitization: involvement of interleukin 10 and transforming growth factor beta. J Exp Med 181, 1887–1892 (1995).

16. D. F. Fiorentino, A. Zlotnik, T. R. Mosmann, M. Howard, A. O’Garra, IL-10 inhibits cytokine production by activated macrophages. J Immunol 147, 3815–3822 (1991).

17. J. Barsig et al., Lipopolysaccharide-induced interleukin-10 in mice: role of endogenous tumor necrosis factor-alpha. Eur J Immunol 25, 2888–2893 (1995).

18. A. M. O’Farrell, Y. Liu, K. W. Moore, A. L. Mui, IL-10 inhibits macrophage activation and proliferation by distinct signaling mechanisms: evidence for Stat3-dependent and - independent pathways. EMBO J 17, 1006–1018 (1998).

19. G. Curtale et al., Negative regulation of Toll-like receptor 4 signaling by IL-10-dependent microRNA-146b. Proc Natl Acad Sci U S A 110, 11499–11504 (2013).

20. G. Grutz, New insights into the molecular mechanism of interleukin-10-mediated immunosuppression. J Leukoc Biol 77, 3–15 (2005).

21. C. Berlato et al., Involvement of suppressor of cytokine signaling-3 as a mediator of the inhibitory effects of IL-10 on lipopolysaccharide-induced macrophage activation. J Immunol 168, 6404–6411 (2002).

22. M. Castellucci et al., IL-10 disrupts the Brd4-docking sites to inhibit LPS-induced CXCL8 and TNF-alpha expression in monocytes: Implications for chronic obstructive pulmonary disease. J Allergy Clin Immunol 136, 781–791 e789 (2015).

23. E. A. Conaway, D. C. de Oliveira, C. M. McInnis, S. B. Snapper, B. H. Horwitz, Inhibition of Inflammatory Gene Transcription by IL-10 Is Associated with Rapid Suppression of Lipopolysaccharide-Induced Enhancer Activation. J Immunol 198, 2906–2915 (2017).

24. D. Kontoyiannis et al., Interleukin-10 targets p38 MAPK to modulate ARE-dependent TNF mRNA translation and limit intestinal pathology. EMBO J 20, 3760–3770 (2001).

25. P. Qasimi et al., Divergent mechanisms utilized by SOCS3 to mediate interleukin-10 inhibition of tumor necrosis factor alpha and nitric oxide production by macrophages. J Biol Chem 281, 6316–6324 (2006).

26. T. Smallie et al., IL-10 inhibits transcription elongation of the human TNF gene in primary macrophages. J Exp Med 207, 2081–2088 (2010).

27. Y. P. Zhu, J. R. Brown, D. Sag, L. Zhang, J. Suttles, Adenosine 5’-monophosphate-activated protein kinase regulates IL-10-mediated anti-inflammatory signaling pathways in macrophages. J Immunol 194, 584–594 (2015).

28. B. Kessler et al., Interleukin 10 inhibits pro-inflammatory cytokine responses and killing of Burkholderia pseudomallei. Sci Rep 7, 42791 (2017).

29. M. Saraiva, P. Vieira, A. O’Garra, Biology and therapeutic potential of interleukin-10. J Exp Med 217, (2020).

30. H. Liu, L. Zeng, Y. Yang, C. Guo, H. Wang, Bcl-3: A Double-Edged Sword in Immune Cells and Inflammation. Front Immunol 13, 847699 (2022).

31. H. Kuwata et al., IL-10-inducible Bcl-3 negatively regulates LPS-induced TNF-alpha production in macrophages. Blood 102, 4123–4129 (2003).

32. Y. Belkaid, O. J. Harrison, Homeostatic Immunity and the Microbiota. Immunity 46, 562–576 (2017).

33. D. S. Shouval et al., Interleukin-10 receptor signaling in innate immune cells regulates mucosal immune tolerance and anti-inflammatory macrophage function. Immunity 40, 706–719 (2014).

34. S. Redpath, P. Ghazal, N. R. Gascoigne, Hijacking and exploitation of IL-10 by intracellular pathogens. Trends Microbiol 9, 86–92 (2001).

35. D. H. Hsu et al., Expression of interleukin-10 activity by Epstein-Barr virus protein BCRF1. Science 250, 830–832 (1990).

36. K. N. Couper, D. G. Blount, E. M. Riley, IL-10: the master regulator of immunity to infection. J Immunol 180, 5771–5777 (2008).

37. T. Fujita, G. P. Nolan, H. C. Liou, M. L. Scott, D. Baltimore, The candidate proto-oncogene bcl-3 encodes a transcriptional coactivator that activates through NF-kappa B p50 homodimers. Genes Dev 7, 1354–1363 (1993).

38. V. Y. Wang et al., Bcl3 Phosphorylation by Akt, Erk2, and IKK Is Required for Its Transcriptional Activity. Mol Cell 67, 484–497 e485 (2017).

39. I. Zanoni et al., CD14 controls the LPS-induced endocytosis of Toll-like receptor 4. Cell 147, 868–880 (2011).

40. A. Gaba et al., Cutting edge: IL-10-mediated tristetraprolin induction is part of a feedback loop that controls macrophage STAT3 activation and cytokine production. J Immunol 189, 2089–2093 (2012).

41. S. Mukhopadhyay et al., Loss of IL-10 signaling in macrophages limits bacterial killing driven by prostaglandin E2. J Exp Med 217, (2020).

42. W. K. E. Ip, N. Hoshi, D. S. Shouval, S. Snapper, R. Medzhitov, Anti-inflammatory effect of IL-10 mediated by metabolic reprogramming of macrophages. Science 356, 513–519 (2017).

43. Y. Guan et al., Cytohesin-4 Upregulation in Glioma-Associated M2 Macrophages Is Correlated with Pyroptosis and Poor Prognosis. J Mol Neurosci 73, 143–158 (2023).

44. J. Thibodeau et al., Interleukin-10-induced MARCH1 mediates intracellular sequestration of MHC class II in monocytes. Eur J Immunol 38, 1225–1230 (2008).

45. K. Gabunia et al., IL-19 Halts Progression of Atherosclerotic Plaque, Polarizes, and Increases Cholesterol Uptake and Efflux in Macrophages. Am J Pathol 186, 1361–1374 (2016).

46. A. F. Alexander, I. Kelsey, H. Forbes, K. Miller-Jensen, Single-cell secretion analysis reveals a dual role for IL-10 in restraining and resolving the TLR4-induced inflammatory response. Cell Rep 36, 109728 (2021).

47. B. M. Tiemeijer, S. Heester, A. Y. W. Sturtewagen, A. Smits, J. Tel, Single-cell analysis reveals TLR-induced macrophage heterogeneity and quorum sensing dictate population wide anti-inflammatory feedback in response to LPS. Front Immunol 14, 1135223 (2023).

48. C. Tudor et al., The p38 MAPK pathway inhibits tristetraprolin-directed decay of interleukin-10 and pro-inflammatory mediator mRNAs in murine macrophages. FEBS Lett 583, 1933–1938 (2009).

49. P. E. Collins, P. A. Kiely, R. J. Carmody, Inhibition of transcription by B cell Leukemia 3 (Bcl-3) protein requires interaction with nuclear factor kappaB (NF-kappaB) p50. J Biol Chem 289, 7059–7067 (2014).

50. X. Guo, A. Adelaja, A. Singh, R. Wollman, A. Hoffmann, Modeling heterogeneous signaling dynamics of macrophages reveals principles of information transmission in stimulus responses. Nat Commun 16, 5986 (2025).

51. C. Zhao, T. X. Medeiros, R. J. Sove, B. H. Annex, A. S. Popel, A data-driven computational model enables integrative and mechanistic characterization of dynamic macrophage polarization. iScience 24, 102112 (2021).

52. G. C. An, J. R. Faeder, Detailed qualitative dynamic knowledge representation using a BioNetGen model of TLR-4 signaling and preconditioning. Math Biosci 217, 53–63 (2009).

53. J. D. Day, D. M. Metes, Y. Vodovotz, Mathematical Modeling of Early Cellular Innate and Adaptive Immune Responses to Ischemia/Reperfusion Injury and Solid Organ Allotransplantation. Front Immunol 6, 484 (2015).

54. R. L. Lopes, T. J. Borges, R. F. Zanin, C. Bonorino, IL-10 is required for polarization of macrophages to M2-like phenotype by mycobacterial DnaK (heat shock protein 70). Cytokine 85, 123–129 (2016).

55. M. D. B. McKay, R.J.; Conover, A Comparison of Three Methods for Selecting Values of Input Variables in the Analysis of Output from a Computer Code. Technometrics 21, 7 (1979).

56. M. W. Pfaffl, A new mathematical model for relative quantification in real-time RT-PCR. Nucleic Acids Res 29, e45 (2001).

57. G. Rousselet, Chromatin Immunoprecipitation in Macrophages. Methods Mol Biol 1784, 177–186 (2018).

58. M. Riemann, R. Endres, S. Liptay, K. Pfeffer, R. M. Schmid, The IkappaB protein Bcl-3 negatively regulates transcription of the IL-10 gene in macrophages. J Immunol 175, 3560–3568 (2005).

59. S. Horber et al., The Atypical Inhibitor of NF-kappaB, IkappaBzeta, Controls Macrophage Interleukin-10 Expression. J Biol Chem 291, 12851–12861 (2016).

